# Dentate Gyrus Norepinephrine Ramping Facilitates Aversive Contextual Processing

**DOI:** 10.1101/2024.10.31.621389

**Authors:** Eric T. Zhang, Grace S. Saglimbeni, Jiesi Feng, Yulong Li, Michael R. Bruchas

## Abstract

Dysregulation in aversive contextual processing is believed to affect several forms of psychopathology, including post-traumatic stress disorder (PTSD). The dentate gyrus (DG) is an important brain region in contextual discrimination and disambiguation of new experiences from prior memories. The DG also receives dense projections from the locus coeruleus (LC), the primary source of norepinephrine (NE) in the mammalian brain, which is active during stressful events. However, how noradrenergic dynamics impact DG-dependent function during contextual discrimination and pattern separation remains unclear. Here, we report that aversive contextual processing in mice is linked to linear elevations in tonic norepinephrine release dynamics within the DG and report that this engagement of prolonged norepinephrine release is sufficient to produce contextual disambiguation, even in the absence of a salient aversive stimulus. These findings suggest that spatiotemporal ramping characteristics of LC-NE release in the DG during stress likely serve an important role in driving contextual processing.

## 1 INTRODUCTION

On any given day, humans must consistently engage in contextual learning processes such as pattern separation and contextual discrimination, as we experience new episodic events and compare novel information to previous experiences. These processes are critical for homeostatic cognitive and affective behavior while deficits in this functioning are thought to underlie multiple neuropsychiatric disease states including post-traumatic stress disorder (PTSD) and schizophrenia ^1,2,3^. These disorders are characterized by a dysregulation of pattern separation, and individuals with these disorders often experience elevated levels of fear and anxiety resulting from failures in contextual processing between incoming episodic information and existing memories, hypervigilance, and exaggerated threat detection ^4,5,6^. Current therapeutic strategies to treat the resulting disorders are limited, although adrenergic receptor antagonists have been recently used with some positive effects ^7,8^. Further research implicates the need for a deeper understanding of the neuromodulatory neural mechanisms behind contextual processing, to open translational opportunities for therapeutic development. The hippocampus is a critical neural structure which mediates learning and memory and is involved in processing of incoming environmental information ^9,10,11,12,13^. Moreover, the dentate gyrus (DG), a subregion of the hippocampus, has been closely linked to contextual learning and pattern separation, wherein information from existing memories is compared to new incoming information ^14,15,16,17^. Additionally, the DG has been shown to be highly modulated by stress and aversive stimuli ^18,19,20^. However, the mechanisms for this remain unresolved. Specifically, how do disrupted memory retrieval and failures in pattern separation give rise to contextual generalization in response to aversive stimuli?

One potential source of modulation for the DG is the locus coeruleus (LC), which provides dense innervations to the DG ^21,22^. The LC, also known as the ‘blue spot’, is a small region located in the brain stem that supplies the mammalian brain with norepinephrine (NE) and has been implicated in regulation of anxiety ^23,24,25,26^. LC-NE tonic activity increases in response to stressful events and is necessary and sufficient for aversive behavior, leading to selective modulation of several brain structures, including the DG ^27,28,29,30,31^. However, it is not known how LC-NE dynamics modulate the DG in memory related disorders, nor what the temporal release properties of NE are in this behavior.

Here, we conducted an integrated neuromodulatory circuit analysis approach, combining optogenetic tools, in vivo fiber photometry, and a NE-selective biosensor with a well-established Pavlovian contextual fear discrimination (CFD) paradigm in which animals were placed in two similar, but different, contexts daily ^12,17,32^. Elucidating the spatiotemporal dynamics of LC-NE modulation of DG-dependent aversive contextual processing provides critical information towards efforts to combat dysregulation of contextual disambiguation in mental health disorders such as PTSD and schizophrenia.

## 2 RESULTS

### Linear elevations of tonic NE release within the DG occur during successful contextual discrimination of an aversive context

To elucidate the spatiotemporal dynamics of NE in the DG during behavior, we first injected novel biosensor GRAB_NE_, a GPCR activation-based NE sensor with high specificity for NE, into the DG of Dbh-cre mice (Fig. 1a-b) ^33^. We then implanted optical fibers above the injection sites for photometry recordings during behavioral tasks including tail lift, foot shock, and optogenetic stimulation (Fig. 1c). To determine expression and efficacy of the GRAB_NE_ biosensor in each animal to be examined, mice first underwent tail lift and foot shock testing for NE release calibration across animals. During tail lift, we observed a significant increase in GRAB_NE_ fluorescence during the tail lift compared to YFP controls (Fig. 1d-e). This signal began to rise significantly when the animal was first picked up by the tail, increased continuously while the animal was suspended for 20 seconds, and decreased back to baseline after the animal was returned to the ground. This outcome matches the dynamics recorded in the lateral hypothalamus during initial development and characterization of GRAB_NE_ ^33^. During foot shock, we also observed a significant increase in GRAB_NE_ fluorescence compared to YFP controls (Fig 1f-g). This signal increased sharply during the 2 second shock and decreased rapidly back to baseline following the termination of the shock. These results indicate successful calibration of GRAB_NE_ for detecting spatiotemporal dynamics of endogenous NE release from LC to DG during stressful stimuli, allowing for potential use in our contextual fear discrimination task.

**FIGURE 1.**
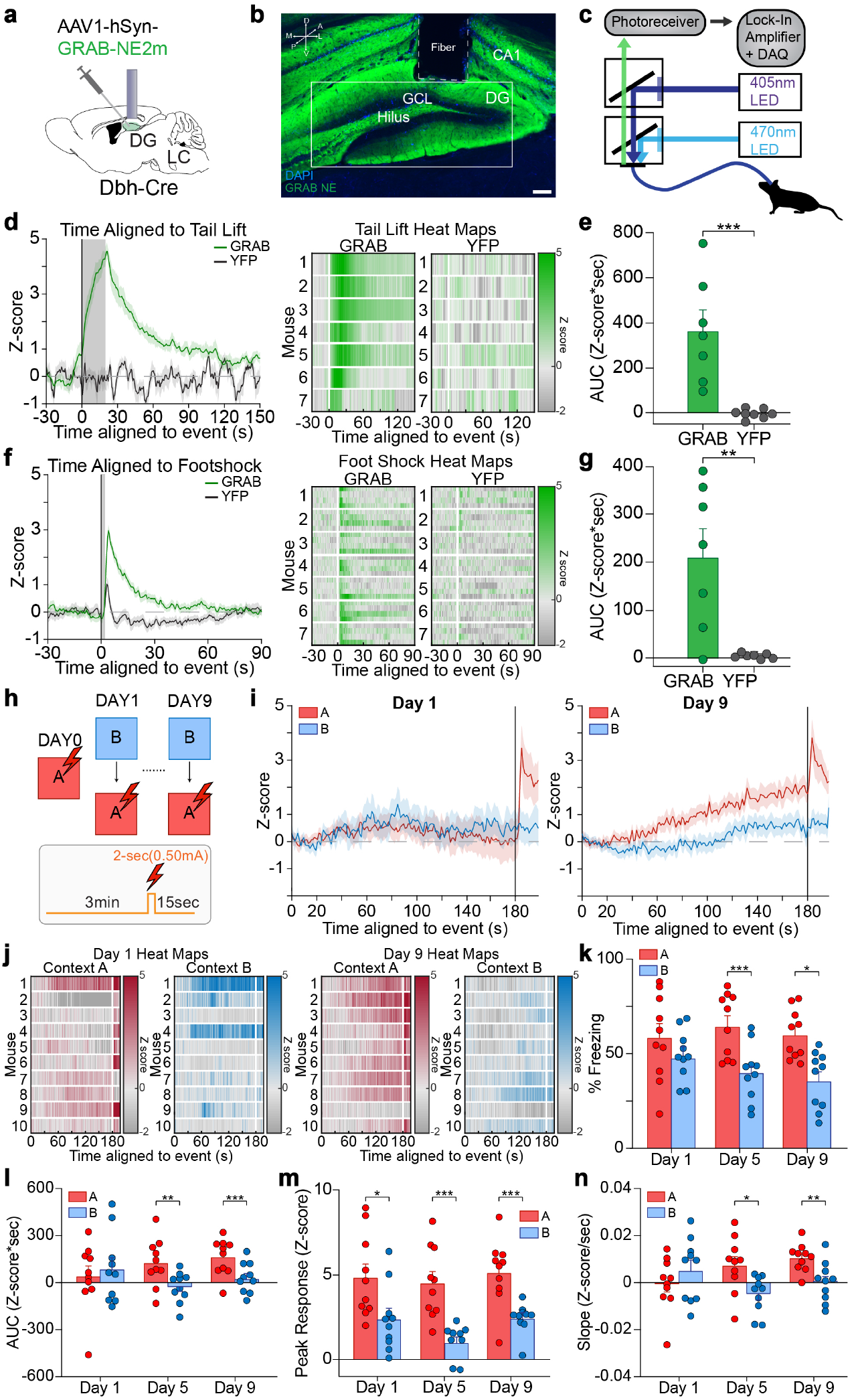
Linear elevations in tonic NE release within the DG occur during successful contextual discrimination of an aversive context. (a) Schematic of experimental approach depicts infection of dentate gyrus with GRAB_NE_, and optical fiber implanted above the DG. (b) Representative image depicting expression of GRAB_NE_ in DG driven by hSyn promoter and location of fiber implant (n = 7 mice). Scale bar = 100 µm. (c) Schematic of fiber photometry setup. (d) Averaged trace of GRAB_NE_ and YFP control fluorescence (left) and individual heat maps of GRAB_NE_ and YFP fluorescence for 20 second tail lift test (right). (e) Area under the curve analysis for GRAB_NE_ and YFP control fluorescence for tail lift test (n = 7 mice). (f) Averaged trace of GRAB_NE_ and YFP control (left) and individual heat maps of GRAB_NE_ and YFP fluorescence for 2 second foot shock (right). (g) Area under the curve analysis for GRAB_NE_ and YFP control fluorescence for 2 second foot shock (n = 7 mice). (h) Schematic depicting CFD task. Animals received a shock in context A (red), but not in context B (blue) for nine days. (i) Averaged traces of GRAB_NE_ fluorescence in context A (red) and context B (blue) for day 1 (left), and day 9 (right). (j) Individual heat maps for context A (red) and context B (blue) for days 1 (left) and 9 (right) of training. (k) Mice successfully discriminated between context A (unsafe), and context B (safe) on the fifth and ninth days of training, but not on the first day of training (n = 10 mice). (l) Area under the curve analysis of GRAB_NE_ fluorescence levels in context A and context B on the first, fifth, and ninth days of training (n = 10 mice). (m) Peak response analysis of GRAB_NE_ fluorescence levels in context A and context B on the first, fifth, and ninth days of training (n = 10 mice). (n) Average slope in context A (red) and context B (blue) on the first, fifth, and ninth days of training (n = 10 mice). All data are mean ± SEM. ^*^*p* < 0.05, ^**^*p* < 0.01, ^***^*p* < 0.005.

We then injected GRAB_NE_ into the DG with optical fibers above the injection site of Dbh-cre mice, which then underwent an established contextual fear discrimination task (Fig. 1h) ^12,17,32^. On day 1, mice displayed no significant differences in NE dynamics during the first 180 seconds of each context, while on days 5 and 9 there was sustained release of NE through the first 180 seconds in context A, while no such elevated release was observed in the safe context B (Fig. 1i-j). These mice also showed robust phasic increases in NE release in response to the 2 second foot shock on both day 1 and day 9 of training. Additionally, while on day 1 there were no significant differences in freezing

We next sought to understand how NE dynamics persisted foll-in context A compared to context B, on day 5 and day 9 there was a significantly increased amount of freezing in context A as compared to context B, indicating that mice successfully discriminated between the unsafe and safe contexts following training (Fig. 1k). To quantify the amount of NE released in both context A and context B, a total rate of change or “area under the curve (AUC)” was calculated, taking the AUC for each mouse in the first 180 seconds of both context A and context B. The amount of NE released in context A was significantly higher than in context B during the middle and late stages of training, while there was no significant difference in AUC between the two contexts in the early stages of training (Fig. 1l). These findings indicate that as mice were increasingly able to distinguish between the safe and unsafe contexts, there was a corresponding increase in NE release in the unsafe context compared to the safe context. To quantify the correlation between these two factors, we calculated a ratio between the proportion of freezing behavior in context A (unsafe) compared to context B (safe), and the change in AUC (ΔAUC) in context A (unsafe) compared to context B (safe). We used linear regression and Pearson correlation to assess the link between these parameters for both the beginning and end of training (Extended Data Fig. 1a). While there is little to no significant relationship between the two variables at the beginning of training, by the end of the task this shifts to a significant fit between freezing ratio and ΔAUC in the two contexts. This indicates that mice that were better able to disambiguate between context A and context B also experienced higher amounts of prolonged NE release in the DG during the CFD task while in the unsafe context. Additionally, we calculated the peak response during the last 17 seconds in each context, to quantify the response to foot shock in context A and saw significantly higher peak response in context A compared to context B, indicating increased NE release in response to the aversive stimulus (Fig. 1m). However, when we used Pearson correlation to assess potential links between the difference in peak response in context A compared to context B with freezing ratio, there were no significant correlations between these two factors (Extended Data Fig. 1b). Additionally, we calculated the slope of the NE dynamics, finding significant differences in slope between context A and context B during the middle and end stages of training, but not the beginning (Fig. 1n). We also tested mice injected with eYFP, in order to ensure that the increases seen during our CFD were in fact increases in NE release, rather than changes in fluorescence levels (Extended Data Fig. 1c-d). While these mice were able to successfully discriminate between context A and context B by the end of training, they did not display any differences in NE release between context A and context B by the end of training, both during the first 180 seconds of the trial and in response to the foot shock (Extended Data Fig. 1e-h). When examining potential links between freezing ratio and AUC and between freezing ratio and peak response via Pearson correlation, there were no significant correlations between these factors (Extended Data Fig. 1i-j). These results suggest that the differences observed during contextual fear discrimination were in fact due to prolonged NE release in the DG, detected by GRAB_NE_. owing successful discrimination in the CFD task and removal of the aversive stimuli, by observing NE signaling during a modified CFD extinction task (Extended Data Fig. 2a-b). In this task, following completion of the standard CFD task, mice underwent two additional days of training in which they were exposed to context B, then context A, for 15 minutes, with no shock in either context. In this task, mice successfully discriminated between context A and context B on both day 5 and day 9 of training, demonstrating successful learning, and continue to display significantly increased freezing in context A compared to context B on Day 10, while on Day 11 these mice no longer show significant differences in freezing between the two contexts. (Extended Data Fig. 2c). On day 1, mice displayed no significant differences in NE dynamics during the first 180 seconds of each context, while on days 5 and 9 there was sustained release of NE through the first 180 seconds in context A, while no such elevated release was observed in the safe context B (Extended Data Fig. 2d-e). These mice also showed robust phasic increases in NE release in response to the 2 second foot shock on both day 1 and day 9 of training. On day 10 of training, mice continue to display increased NE release in context A compared to context B, with NE levels remaining elevated through the entirety of the 15-minute trial, while on day 11 there are no significant differences in NE levels between the two contexts (Extended Data Fig. 2f-g). To quantify the total amount of NE released in both context A and context B over time, an area under the curve (AUC) analysis was used, taking the AUC for each mouse in the first 180 seconds of both context A and context B. The amount of NE released in context A was significantly higher than in context B during days 5, 9, and 10, while there was no significant difference in AUC between the two contexts on days 1 and 11 (Extended Data Fig. 2h). Additionally, we calculated the peak response between 180 and 197 seconds in each context, to quantify the response to foot shock in context A and saw significantly higher peak response in context A compared to context B on days 1, 5, and 9, indicating increased NE release in response to the aversive stimulus (Extended Data Fig. 2i). There was also significantly increased peak response in context A compared to context B on day 10 but not on day 11, indicating the strength of the sustained NE release in context A compared to context B on day 10. We also calculated the slope of the NE dynamics, finding significant differences in slope between context A and context B during days 9 and 10 of training, but not days 1, 5, and 11 (Extended Data Fig. 2j). These results suggest that prolonged NE release in the DG persists following contextual discrimination even in the absence of the aversive stimuli, and that the termination of this NE ramping occurs in association with the end of contextual learning.

Next, we aimed to elucidate potential differences in NE ramping due to repeated exposure to aversive stimuli and one extremely aversive stimulus. This was done through a modified CFD experiment that took place over two days, with increased foot shock intensity (Extended Data Fig. 3a-b). In this modified CFD task, mice successfully discriminated between context A and context B on the second day and displayed increased NE dynamics on day 2 in the unsafe context (Extended Data Fig. 3c-e). When these dynamics were quantified via an area under the curve (AUC) analysis, we observed a significantly increased amount of NE in context A compared to context B on day 2 during the first 180 seconds, as well as a significantly increased amount of NE in context A compared to context B in response to the foot shock (Extended Data Fig. 3f-g). We then used Pearson correlation to assess potential links between the difference in AUC and peak response in context A compared to context B with freezing ratio, find that there were no significant correlations on day 1, but that there was a significant correlation between ΔAUC and freezing ration on day 9 (Extended Data Fig. 3h-i). Lastly, we calculated the slope of the NE dynamics, finding no significant differences in slope between context A and context B during either day of training (Extended Data Fig. 3j). These findings suggest similar NE dynamics in the DG during varying types of contextual fear learning.

Given that the hippocampus also likely receives dopamine as a byproduct of NE synthesis, it is possible that this neuromodulator may also be important in aversive contextual learning ^34,35^. To this end, we repeated the CFD experiment with novel biosensor GRAB_DA_, a GPCR activation-based DA sensor with high specificity for DA, into the DG of DAT-cre mice (Extended Data Fig. 4a-b). On day 1 there were no significant differences in freezing in context A compared to context B, however on day 5 and day 9 there was a significantly increased amount of freezing in context A as compared to context B, indicating that mice successfully discriminated between the unsafe and safe contexts following training (Extended Data Fig. 4c). We also observed elevated dopamine signaling in both context A and context B by the end of training, as well as elevated responses in response to the shock (Extended Data Fig. 4e-h). Additionally, when we used Pearson correlation to assess potential links between the differences in AUC in context A compared to context B with freezing ratio, there were no significant correlations between these two factors (Extended Data Fig. 4i). There was however a significant correlation on day 9 when using Pearson correlation to assess potential links between the differences in peak response in context A compared to context B with freezing ratio (Extended Data Fig. 4j). Lastly, after calculating the slope of the DA dynamics, we found no significant differences between context A and context B during the early, middle, and late stages of training (Extended Data Fig. 4k). These results indicate that while DA is released in the DG during CFD, contextual discrimination as it relates to aversive content specifically is likely to not be mediated by DA in this paradigm. However, the role of dopamine in CFD specifically requires substantial further evaluation, additional sensor selectivity types, and its own complete study.

### Induction of elevations in DG NE dynamics is sufficient to cause contextual disambiguation in the absence of a salient aversive stimulus

Following characterization of GRAB_NE_ in the DG during endogenous NE release, we next characterized GRAB_NE_ dynamics in response to evoked NE release. We injected GRAB_NE_ into the DG, and Cre-recombinase dependent viral vectors containing the red shifted light-sensitive cation channel ChrimsonR into the LC of Dbh-cre mice (Fig. 2a-b). We then implanted optical fibers above the DG for photometry recordings during contextual discrimination and began testing stimulation parameters to tune photo-stimulation to closely match endogenous NE dynamics seen in CFD (Fig. 2c). 20 Hz pulsed photostimulation for 20 seconds was delivered above the DG, during which we observed a significant increase in GRAB_NE_ fluorescence during stimulation compared to YFP controls (Fig. 2d-e). This signal then decreased back to baseline following termination of the stimulation parameters. In a new series of experiments, we also delivered 1, 3, 5, and 10 Hz pulsed photostimulation for 20 seconds in separate trials and observed increasing levels of GRAB_NE_¬ fluorescence with increasing levels of pulsed photostimulation of LC-DG terminals, suggesting that we are able to discreetly control evoked release of NE from LC to DG (Fig. 2f-h). These results establish successful characterization of GRAB_NE_ for detecting spatiotemporal dynamics of evoked NE release from LC to DG, providing an optogenetic titration and calibration template for use in our contextual fear discrimination task.

**FIGURE 2.**
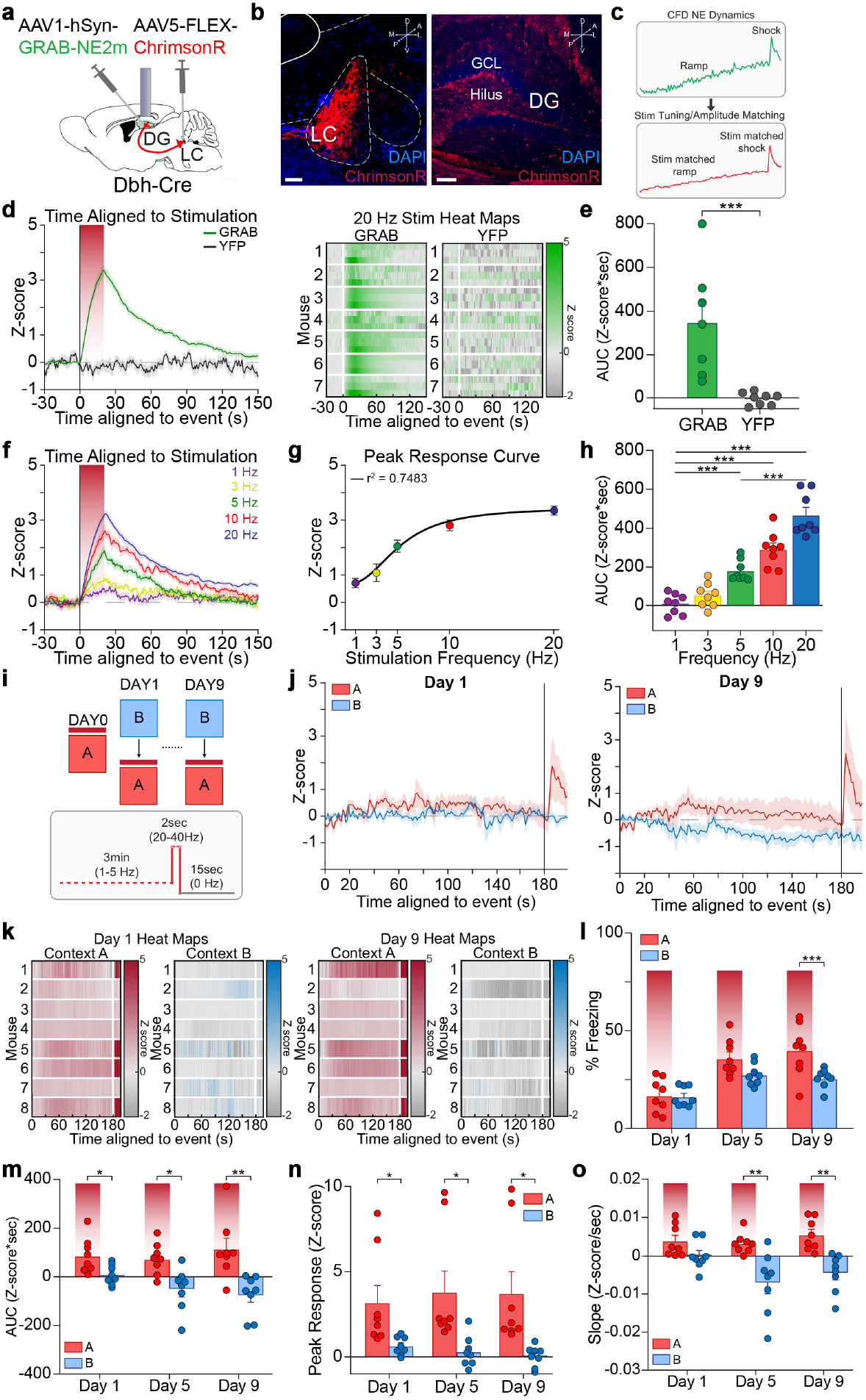
Optogenetic control of linear elevations in DG NE dynamics is sufficient to cause contextual disambiguation in the absence of a salient aversive stimulus. (a) Schematic of experimental approach depicts infection of DG with GRAB_NE_, and optical fiber implanted above the DG, and infection of LC with ChrimsonR. (b) Representative image depicting expression of ChrimsonR in LC (left) and terminal expression in DG (right) (n = 7 mice). Scale bar = 100 µm. (c) Schematic depicts stimulation tuning and amplitude matching for stimulation parameters comparing to endogenous NE dynamics observed during CFD. (d) Averaged trace of GRAB_NE_ and YFP control fluorescence (left) and individual heat maps of GRAB_NE_ fluorescence for 20 Hz photostimulation for 20 seconds (right). (e) Area under the curve analysis for GRAB_NE_ and YFP control fluorescence for 20 Hz photostimulation for 20 seconds (n = 7 mice). (f) Averaged traces of GRAB_NE_ fluorescence for 1, 3, 5, 10 and 20 Hz photostimulation for 20 seconds. (g) Peak response curve for variable photostimulation (n = 8 mice). (h) Area under the curve analysis for GRAB_NE_ fluorescence for variable photostimulation (n = 8 mice). (i) Schematic depicting CFD task. Animals received stimulation in context A (red), but not in context B (blue) for nine days. (j) Averaged traces of GRAB_NE_ fluorescence in context A (red) and context B (blue) for day 1 (left), and day 9 (right). (k) Individual heat maps for context A (red) and context B (blue) for days 1 (left) and 9 (right) of training. (l) Mice successfully discriminated between context A (unsafe), and context B (safe) on the fifth and ninth days of training, but not on the first day of training (n = 8 mice). (m) Area under the curve analysis of GRAB_NE_ fluorescence levels in context A and context B on the first, fifth, and ninth days of training (n = 8 mice). (n) Peak response analysis of GRAB_NE_ fluorescence levels in context A and context B on the first, fifth, and ninth days of training (n = 8 mice). (o) Average slope in context A (red) and context B (blue) on the first, fifth, and ninth days of training (n = 8 mice). All data are mean ± SEM. ^*^*p* < 0.05, ^**^*p* < 0.01, ^***^*p* < 0.005.

We next examined how this increased and sustained release of NE in the aversive context during CFD would alter discrimination and recognition during contextual processing. To do this, we utilized GRAB_NE_ with an optical fiber implanted in the DG, and injected ChrimsonR into the LC of Dbh-cre mice (Fig. 2a-b). These mice underwent a modified CFD task, in which the foot shock was entirely replaced by a variable photo-stimulation of LC-DG terminals while in context A in order to closely match the NE linear ramping effects induced by shock (Fig. 2c, i). During the first 180 seconds of context A, mice received 1-5 Hz pulsed photostimulation, mimicking tonic NE release dynamics and resulting in sustained NE release over the session. Then, mice received 20-40 Hz pulsed photostimulation for 2 seconds, to replicate the phasic bursting we measured during responses to foot shock. The frequencies of stimulation that mice received were specifically calibrated for each individual mouse to mimic the endogenous NE peak response and dynamics measured during CFD, with stim matched ramping during the first 180 seconds and stim matched foot shock for 2 seconds in context A (Fig. 2j-k). Mice that underwent this modified CFD task displayed no differences in freezing between contexts in early or middle stages of training and displayed significantly increased freezing in context A compared to context B during the late stages of training, even in the absence of a salient aversive stimuli (Fig. 2l). To quantify the total amount of NE released in both context A and context B over time, an area under the curve (AUC) analysis was used, taking the AUC for each mouse in the first 180 seconds of both context A and context B. Stimulation occurred across each day of training, with significantly higher NE levels in the aversive context during early (day 1), middle (day 5) and late (day 9) stages of training, suggesting that the stimulation parameters chosen were sufficient to evoke prolonged NE release in the DG in context A throughout the CFD task (Fig. 2m). These findings demonstrate increased learning over the course of the CFD task, in which mice show they are able to learn how to disambiguate between the two contexts by freezing more in context A, the “unsafe” context due to increased NE release, even in the absence of an aversive stimuli. To quantify whether there was a linear correlation between these two factors, we calculated a ratio between the proportion of freezing behavior in context A (unsafe) compared to context B (safe), and the change in AUC in context A (unsafe) compared to context B (safe). We then used linear regression and Pearson correlation to determine the relationship between these ratios for both the beginning and end of training (Extended Data Fig. 5a). While there is no significant relationship between the two variables at the beginning of training, by the end of the task there is a significant correlation between freezing ratio and ΔAUC within the two contexts. This result indicates that mice receiving higher amounts of prolonged NE release in the “unsafe” context also performed better by the end of training in the CFD task, learning to disambiguate the two contexts in the absence of an aversive stimuli. Additionally, we also quantified the peak response for each mouse in response to the foot shock in context A, with mice showing significantly increased NE release in context A compared to context B at 180 seconds, when the foot shock was administered (Fig. 2n). In order to quantify a potential relationship between freezing and peak response, we use a Pearson correlation analysis between the difference in peak response in context A vs. context B and the freezing differences in context A vs. context B (Extended Data Fig. 5b). Here, no significant correlation was found between freezing and peak response in both the beginning and end of training. Furthermore, we calculated the slope of the NE dynamics, finding significant differences in slope between context A and context B during the middle and end stages of training, but not the beginning (Fig. 2o). During this modified contextual fear paradigm with stimulation matching, in context B, we noticed a decaying signal that was not seen prior. We hypothesized that this decay was due to a lack of salient stimuli during this altered CFD paradigm. Therefore, we conducted another altered CFD experiment in which mice were placed in both context A and B for 197 seconds with no salient stimuli or stimulation (Extended Data Fig. 5c-d). Mice that received no stimulation in either of the two contexts displayed no significant differences in freezing, AUC, slope, and correlation between freezing ratio and ΔAUC over the course of training (Extended Data Fig. 5e-j).Taken together, these results indicate that mice experience increased discrimination between two contexts when NE is increased in a linear manner within the DG in one context, in the absence of a salient aversive stimuli such as foot shock, and that the stronger the tonic elevation in NE release is over the trial, the better the performance of disambiguating both contexts in the CFD task.

### Mimicking endogenous ramping of NE tonic release in the DG is sufficient for contextual disambiguation in the absence of a salient aversive stimulus

We next aimed to isolate the specific spatiotemporal NE release dynamics responsible for causing discrimination between two contexts, in the absence of a salient stimuli. In order to do this, we again utilized GRAB_NE_ with an optical fiber implanted in the DG, and injected ChrimsonR into the LC of Dbh-cre mice for use in fiber photometry (Fig. 3a). We again used a modified CFD task, during which we reproduced the NE release event in response to a foot shock by delivering 20-40 Hz pulsed photostimulation for 2 seconds after 180 seconds in context A (Fig. 3b-c). Here, we were again able to effectively calibrate stimulation to mimic endogenous NE dynamics measured during the foot shock delivered during CFD, with stimulation matched shock (Fig. 3d-e). When these mice were stimulated to reproduce a foot shock NE release event during CFD, across each day of training we observed no significant differences in freezing between the two contexts (Fig. 3f). These mice also displayed no differences in AUC during the first 180 seconds of training, suggesting no endogenous prolonged NE release was occurring (Fig. 3g) due to the accumulation of foot-shock release events as the trial unfolded. We again calculated the ratio between the proportion of freezing behavior in context A (unsafe) compared to context B (safe), and the change in AUC in context A (unsafe) compared to context B (safe). We used linear regression and Pearson correlation to assess the relationship between these ratios for both the beginning and end of training. In this case, no correlative relationship was observed between the freezing ratio and ΔAUC in mice that received only 2 seconds of 20-40 Hz pulsed photostimulation throughout training (Extended Data Fig. 6a-b). Additionally, we also quantified the peak NE release response in each mouse due to high frequency stimulation in context A, with mice showing significantly increased NE release in context A compared to context B at 180 seconds (Fig. 3h). In order to quantify a potential relationship between freezing and peak response, we used a Pearson correlation between the difference in peak response in context A vs. context B and the freezing differences in context A vs. context B (Extended Data Fig. 6c). No significant correlation was found between freezing and peak response at both the beginning and end of training. We again calculated the slope of the NE dynamics, finding no significant differences in slope between context A and context B during the CFD task (Fig. 3i).

**FIGURE 3.**
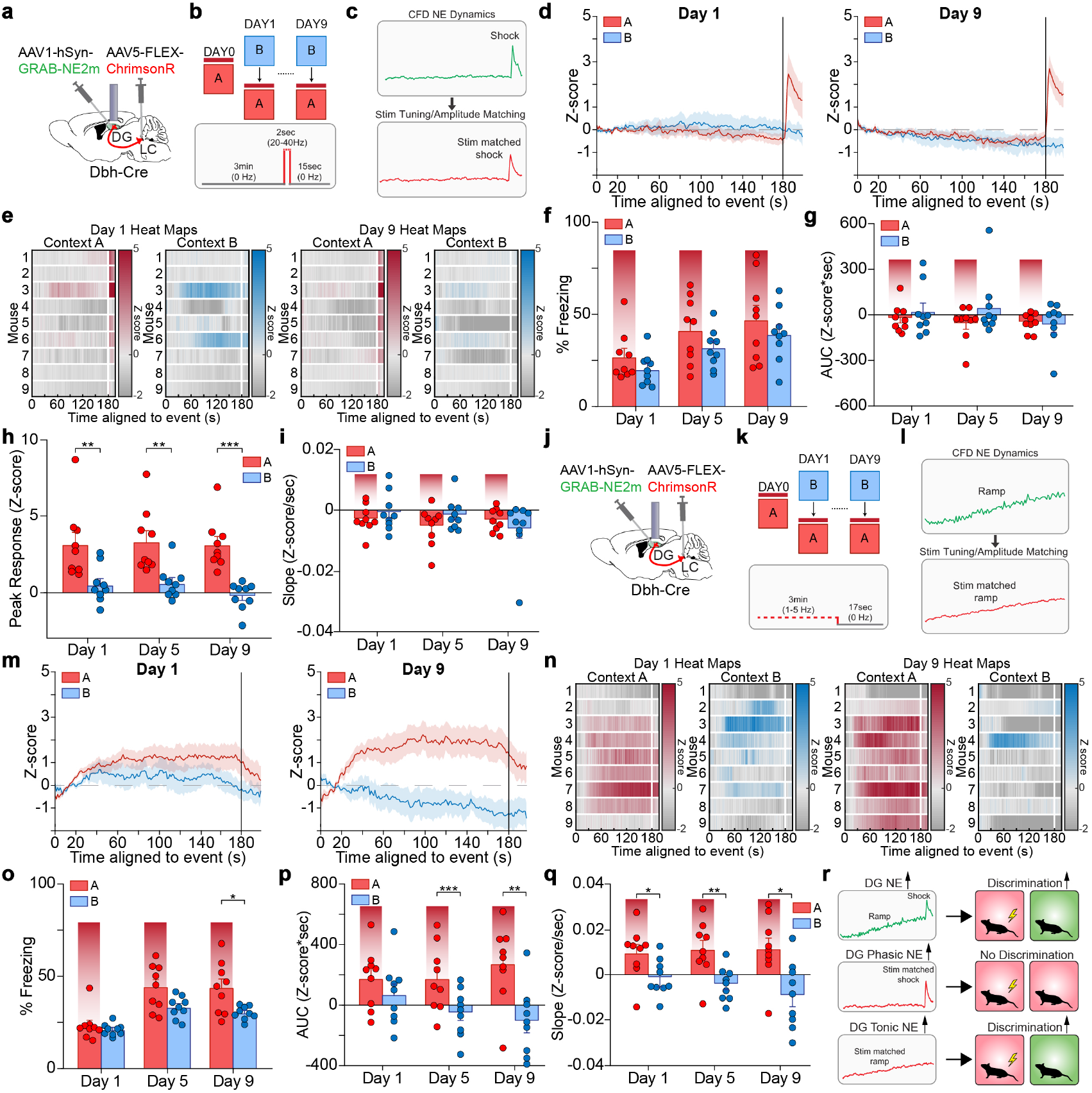
Mimicking endogenous ramping of NE tonic release in the DG is sufficient for contextual disambiguation in the absence of a salient aversive stimulus. (a) Schematic of experimental approach depicts infection of DG with GRAB_NE_, optical fiber implanted above the DG, and infection of LC with ChrimsonR. (b) Schematic depicting CFD task. Animals received stimulation in context A (red), but not in context B (blue) for nine days. (c) Schematic depicts stimulation tuning and amplitude matching for stimulation parameters comparing to endogenous NE dynamics observed during CFD. (d) Averaged traces of GRAB_NE_ fluorescence in context A (red) and context B (blue) for day 1 (left), and day 9 (right). (e) Individual heat maps for context A (red) and context B (blue) for days 1 (left) and 9 (right) of training. (f) Mice were not able to successfully discriminate between context A (unsafe), and context B (safe) any days of training (n = 9 mice). (g) Area under the curve analysis of GRAB_NE_ fluorescence levels in context A and context B on the first, fifth, and ninth days of training (n = 9 mice). (h) Peak response analysis of GRAB_NE_ fluorescence levels in context A and context B on the first, fifth, and ninth days of training (n = 9 mice). (i) Average slope in context A (red) and context B (blue) on the first, fifth, and ninth days of training (n = 9 mice). (j) Schematic of experimental approach depicts infection of DG with GRAB_NE_, optical fiber implanted above the DG, and infection of LC with ChrimsonR. (k) Schematic depicting CFD task. Animals received stimulation in context A (red), but not in context B (blue) for nine days. (l) Schematic depicts stimulation tuning and amplitude matching for stimulation parameters comparing to endogenous NE dynamics observed during CFD. (m) Averaged traces of GRAB_NE_ fluorescence in context A (red) and context B (blue) for day 1 (left), and day 9 (right). (n) Individual heat maps for context A (red) and context B (blue) for days 1 (left) and 9 (right) of training. (o) Mice successfully discriminated between context A (unsafe), and context B (safe) on the fifth and ninth days of training, but not on the first day of training (n = 9 mice). (p) Area under the curve analysis of GRAB_NE_ fluorescence levels in context A and context B on the first, fifth, and ninth days of training (n = 9 mice). (q) Average slope in context A (red) and context B (blue) for early, middle, and late stages of training (n = 9 mice). (r) Schematic depicting major finding that elevation of NE tonic release in the DG is sufficient for contextual disambiguation in the absence of a salient aversive stimulus. All data are mean ± SEM. ^*^*p* < 0.05, ^**^*p* < 0.01, ^***^*p* < 0.005.

Next, we used another modified CFD task, in which the prolonged NE release was mimicked by variable stimulation of LC-DG terminals while in context A to match stimulation to endogenous NE dynamics during CFD (Fig. 3j-l). In this case, we delivered only 1-5 Hz pulsed photostimulation for 180 seconds, without the 2 seconds of high frequency stimulation afterwards, to mimic only the prolonged NE release seen in context A during CFD. This resulted in prolonged tonic NE release dynamics similar to those originally observed during the traditional CFD paradigm (Fig. 3m-n). During this modified CFD paradigm with stimulation matched ramping, mice showed significantly increased freezing in context A compared to context B on day 9 of the CFD task, while no differences in freezing were observed on day 1 or day 5 of the task (Fig. 3o). To quantify the amount of NE released in both context A and context B, an area under the curve (AUC) analysis was used, taking the AUC for each mouse in the first 180 seconds of both context A and context B. Stimulation occurred across each day of training, only in context A, leading to significantly increased NE levels in context A compared to context B across the middle and late stages of training (Fig. 3p). These findings indicate that over the course of training, mice were able to discriminate successfully between context A and context B via the presence of prolonged elevated tonic NE release in context A, even in the absence of a salient aversive stimuli or a foot shock-like evoked phasic NE response. We next calculated a ratio between the proportion of freezing behavior in context A (unsafe) compared to context B (safe), and the change in AUC in context A (unsafe) compared to context B (safe). We used linear regression and Pearson correlation to assess the relationship between these ratios for both the beginning and end of training (Extended Data Fig. 6d-e). While no significant relationship between the two variables was observed at the beginning of training, by the end of the task there was a significant correlation between the freezing ratio and ΔAUC in the two contexts. When we calculated the slope of the NE dynamics, we found significant differences in slope between context A and context B during the beginning, middle and end stages of training (Fig. 3q). These results indicate that linear elevations in tonic NE release within the DG in response to contextual cues is a critical driving force behind contextual discrimination of a safe versus unsafe context (Fig. 3r). In addition, our data suggest that a single robust phasic burst release of NE within the DG as related to the aversive stimuli is not sufficient to produce robust contextual discrimination in response to an aversive stimuli. Taken together, these results establish a critical role for linear tonic increases (ramps) of NE release in the DG as crucial for promoting contextual discrimination.

## 3 DISCUSSION

In the current study, we report that a particular pattern of linear ramping of LC-NE release into the DG results in successful contextual discrimination between a safe and unsafe context in an aversive contextual discrimination task. This is likely through elevated LC-NE neuron tonic firing in response to environmental cues ^36^. We observe sustained ramping of NE release in an aversive context during successful contextual disambiguation in an aversive fear discrimination task and we were able to cause contextual disambiguation using optogenetic mimicry of evoked NE release temporal dynamics. These results further establish the noradrenergic DG interactions as critical for differentiating between new and previous events and indicate that responses to contextual differences are determined from the incoming spatiotemporal dynamics of NE release during an aversive experience.

Previous work has implicated prolonged phasic dopamine signals in striatum as important in signaling proximity and value of reward ^37^. In this study, it was postulated that ramping dopamine signals may provide a continuous estimate of distance and size of reward and maintaining motivation towards that reward. Additionally, further work implicates burst stimulation of ventral tegmental area (VTA) neurons in causing a prolonged increase of dopamine in the nucleus accumbens and prefrontal cortex ^38^. Here however, we demonstrate prolonged ramping norepinephrine release in response to contextual discrimination, in which this NE ramping occurs in the time leading up to delivery of a salient aversive foot shock in the CFD task, during which the animal must rely on contextual cues to disambiguate the unsafe (shock) and safe (no shock) contexts (Fig. 1). In addition, while in prior studies of monoamine ramping the prolonged increases in tone occurred for up to 10 seconds, the NE ramping we report here is exciting because it is quite sustained, lasting for 180 seconds prior to the foot shock stimuli. This suggests that there is an extended time scale function for ramping NE release within the DG, especially during pattern separation, where continuous processing on contextual clues may be necessary for successful discrimination to occur. In addition, we demonstrated prolonged ramping dopamine release in both contexts during contextual discrimination, suggesting a potential role for ramping dopamine during contextual fear learning as well (Extended Data Fig. 4). However, because the ramping takes place in both contexts, this signal appears to be unique from the ramping norepinephrine observed. It is possible that dopamine ramping is responsible for spatial processing, rather than contextual discrimination. Further investigations are needed to more clearly understand how these prolonged neurotransmitter dynamics are mechanistically involved in contextual learning. For example, whether these prolonged ramps of NE release are involved in specific phases of memory such as encoding, consolidation, or extinction. Our results suggest that it will be important to further study the specific methods by which dopamine and norepinephrine regulate contextual discrimination through reuptake, receptor selective binding and signaling, and to better isolate the roles of these neurotransmitters in DG function ^39^.

In this study, we opted to use both male and female mice in the CFD task. Existing literature supports potential sex differences in aversive contextual processing in both rodents and humans, with females more likely to present with symptoms and be diagnosed with PTSD ^40^. In addition, female rats outperformed male rats in a contextual fear discrimination task, while male rats outperformed female rats in a Morris water maze spatial learning task ^41,42^. However, in our CFD task, no differences in contextual discrimination performance were observed between male and female mice (Extended Data Fig. 1l). This may be due to differences in the contextual fear conditioning procedure used, as well as differences in the difficulty of the CFD task. Because sex is a prominent risk factor for development of PTSD in human trauma-exposed adults, further understanding of the potential sex differences in LC-DG mediated pattern separation may be important in development of therapeutic strategies for treatment of PTSD and PTSD-like symptoms ^43,44^.

Throughout this study, we opted to continue use of an established contextual fear conditioning paradigm, as demonstrated in other previous work ^12,17,32^. In this CFD task, animals undergo 9 days of training in which context B is presented before context A. In our testing, when context A and B presentations were alternated, this created a task that was difficult to the point that mice were unable to successfully discriminate between the safe and unsafe contexts ^17^. Thus, maintaining the same order of contextual presentation is necessary to create a baseline in which the animals are able to discriminate between the contexts, so that norepinephrine signaling could be observed during this successful discrimination. Furthermore, during this task we opted to continue the use of cinnamon and peppermint as odorants for the unsafe and safe contexts, respectively. While peppermint odor has been shown to be aversive at higher concentrations, this would not affect our experimental findings, as we used a low dose of peppermint odor in the safe context, while mice showed higher freezing levels in the unsafe context, which contained cinnamon odor instead. Therefore, the discrepancy in freezing levels is not due to the use of a potentially aversive odorant.

We also conducted several modified CFD experiments in this work, including an extinction paradigm and a shortened, heightened aversion paradigm (Extended Data Figs. 2 and 3). During the extinction paradigm, we observed prolonged NE release in context A on day 10 even without presentation of the foot shock, with this prolonged released remaining elevated throughout the entirety of the 15-minute trial. This was accompanied by increased freezing in context A compared to context B. However, on day 11, this ramping no longer took place, and the animals learned to disassociate context A from the foot shock, as there were no significant differences in freezing between context A and context B. These findings further support that as mice undergo contextual learning, there is a corresponding change in prolonged NE dynamics in the DG to inform their behavior. Meanwhile, with the shortened 2-day CFD task, we aimed to understand potential differences in NE dynamics in response to one extremely salient stimulus, as opposed to repeated exposures. In this case, we found that animals do still undergo NE ramping in context A following successful learning of the CFD task, although the change was not as pronounced as during our standard 10-day CFD task. This experiment suggests that similar but not identical mechanisms exist for varying types of contextual learning, which could help inform our treatment of those suffering from PTSD, as there may be differences in development of PTSD following repeated exposure to traumatic events compared to exposure to a singular highly traumatic event. Thus, further research must be done regarding the mechanisms by which norepinephrine regulates contextual discrimination and to better isolate the roles of norepinephrine in DG function.

Additionally, during the modified CFD task used here wherein mice received one of the four different stimulation parameters, there appeared to be gradual decreases in NE tone (quantified by AUC analysis) over the course of training in context B, where no stimulation occurred (Fig. 2, Fig. 3, Extended Data Fig. 5). This is likely due to the established role for LC-NE in novelty encoding as well as contextual learning ^45,46^. We postulate that during this contextual fear discrimination task, and in the absence of an aversive stimuli, mice undergo increased NE release in both contexts during the initial stages of training, as they experience these novel contexts for the first time. However, as these mice become habituated to the contexts, and without any salient cues to respond to, this novelty encoding likely decreases, leading to decreased amounts of NE in the DG during the CFD task, which is then overcome in context A via increased evoked NE release due to photostimulation of LC-DG terminals. There also appear to be high variances in basal anxiety states between distinct cohorts of mice, as seen in the freezing levels during the initial stages of contextual fear discrimination. This is known in the field, and we do our best to control for these differences with animal handling, and reverse light cycle housing. However, these variances were also observed in our previous efforts using this discrimination task, and differences in initial freezing during the first day of training do not appear to impact overall performance in the task by the end of training ^17^.

Overall, our findings presented here provide us with a better understanding of the spatiotemporal properties of norepinephrine release and its time-locked receptor-mediated signaling actions during aversive contextual discrimination. We demonstrated here that increases in tonic ramping of NE release occurs within the DG during successful contextual discrimination. Additionally, mimicking this ramping effect through photoactivation of LC-DG terminals is sufficient to drive pattern separation and contextual disambiguation in the CFD task, even in the absence of a salient aversive stimuli such as foot shock resolving when, where and how NE release in hippocampus, specifically the DG, influences behavior. Understanding the spatiotemporal mechanisms by which monoamine neuromodulators coordinate aversive processing will aid in the development of novel therapeutic strategies for targeting psychiatric disorders characterized by generalization, including PTSD and anxiety.

## 4 MATERIALS AND METHODS

### Animals

Adult (20-35 g) Dbh-Cre mice, DAT-IRES-cre mice, and Cre (-) littermate control mice were used for projection mapping and all *in vivo* experiments after backcrossing to C57BL/6J mice for at least 10 generations. Mice were group housed, given access to food pellets and water *ad libitum*, and maintained on a 12:12-hour reverse light/dark cycle (lights on at 8:00 p.m.). Animals were held in a sound attenuated holding room facility in the lab starting at least one week prior to surgery, as well as post-surgery and throughout the duration of behavioral assays to minimize stress from transportation and disruption from foot traffic. All mice were handled and, where appropriate, connected to fiber optics two times a day for one week prior to behavioral experimental testing. All experimental procedures were approved by the Animal Care and Use Committee of University of Washington and conformed to UW National Institutes of Health guidelines.

### Stereotaxic Surgery

After acclimatizing to the holding facility for 7–9 days, the animals were anesthetized in an induction chamber (4% isoflurane, Piramal Healthcare, Maharashtra, India) and mounted on a stereotaxic frame under sterile conditions (Kopf Instruments, Model 1900) where they were maintained at 1–2% isoflurane for the duration of the surgery. For *in vivo* fiber photometry experiments, adult mice were injected bilaterally with 800 nl in each side of AAV5/Syn-FLEX-ChrimsonR-td-Tomato (Addgene) into the LC (AP: -5.45 mm, ML: ±1.25 mm, DV: -3.8 mm) using a Hamilton syringe with a beveled needle and injected unilaterally with 400 nl of AAV1-hsyn-GRAB-NE2m or AAV9-hsyn-GRAB-DA2m (Yulong Li Lab) into the DG (AP: -2.15 mm, ML: +1.4 mm, DV: -2.1 mm) using a Hamilton syringe with a blunted needle. Transgenic controls were injected bilaterally with 800 nl of AAV5/Syn-FLEX-ChrimsonR-td-Tomato into the LC (AP: -5.45 mm, ML: ±1.25 mm, DV: -3.8 mm) using a Hamilton syringe with a beveled needle and injected unilaterally with 400 nl of AAV5-Ef1a-DIO-EYFP (Addgene) into the DG using a Hamilton syringe with a blunted needle. Mice then received intracranial fiber photometry implants in the DG (AP: -2.15 mm, ML: +1.4 mm, DV: -1.7 mm), which were secured using MetaBond (C & B Metabond). Mice were allowed to recover for at least six weeks following infusion of virus and intracranial implant prior to further behavioral testing or perfusion for projection mapping to ensure optimal viral expression. Mice were perfused at the conclusion of behavior to ensure optimal viral expression and optical implant placement location (Extended Data Fig. 6f).

### Tissue Collection and Immunohistochemistry

After the conclusion of behavioral testing, mice were anesthetized with sodium pentobarbital and transcardially perfused with ice-cold PBS, followed by 4% phosphate-buffered paraformaldehyde. Brains were removed, postfixed overnight in 4% paraformaldehyde, and then saturated in 30% phosphate-buffered sucrose for 2-4 days at 4°C. Brains were sectioned at 30 mM on a microtome and stored in a 0.01M phosphate buffer at 4°C prior to immunohistochemistry and tracing experiments. For behavioral cohorts, viral expression and optical fiber placements were confirmed before inclusion in the presented datasets. Viral expression and implant placements were verified using fluorescence and confocal (Leica Microsystems) microscopy. Images were produced with 10X, 20X, 63X objective and analyzed using ImageJ software (NIH) and Leica Application Suite Advanced Fluorescence software.

### Fear Conditioning

Fear conditioning took place in Med-Associates conditioning chambers that consisted of one clear plexiglass wall, three aluminum walls, and a stainless-steel grid as a floor. The training chamber was housed in a sound attenuated cubicle. The conditioning chambers could be configured into two distinct contexts: A and B. Context A was rectangular, with floors made of stainless-steel rods (2 mm diameter, spaced 5 mm apart), walls of aluminum and acrylic, and cinnamon extract scent, and was cleaned with 70% ethanol between runs. Context B differed from context A in that it had a white acrylic floor, and a wall decorated with a black and white striped pattern. Context B was cleaned with multipurpose surface cleaner (Method, UPC: 817939000106) between runs, and scented with peppermint extract. All sessions were recorded from the side using a digital camera and were scored for freezing by an investigator blinded to the genotype and experimental conditions of the animal.

For *in vivo* fiber photometry during behavior tests, a rotating optical commutator (Doric) was positioned on top of the training chamber and connected to a fiber photometry recording rig (ThorLabs). Fibers were attached to the implants on the mouse for every training session.

### Contextual Fear Discrimination (CFD)

Mice were handled for 5 minutes per day for a week prior to training. Contextual fear discrimination training took place in the apparatus described above. During a standard CFD experiment, in context A (unsafe context), animals were placed in the conditioning chamber and allowed to freely explore for 180 seconds, after which they received a single 2 second foot shock of 0.50 mA. Mice were taken out 15 seconds after termination of the foot shock and returned to their home cage. In context B (safe context), mice were allowed to freely explore the context for 197 seconds, the same amount of total time that they were in context A, and then returned to their home cage. In this context, no foot shock was delivered. On the first day (Day 0) of CFD training, animals were placed in context A and received a foot shock. The next day (Day 1), mice were run first in context B with no foot shock, then in context A with a foot shock. There was a minimum 3-hour period between exposure to context A and context B. This training protocol was repeated daily for 9 days. All freezing in both contexts was assessed for the first 180 seconds, and the last 17 seconds were not included in the behavioral analysis. Freezing was done manually by an investigator in a double-blind fashion to avoid bias in scoring. Mice only underwent one CFD experiment.

For the modified extinction CFD experiment, mice followed the standard CFD experiment described above through day 9 (Extended Data Fig. 2). Then, on days 10 and 11, mice were placed into the conditioning chamber and allowed to freely explore for 15 minutes, after which they were taken out and returned to their home cage. They were run first in context B with no foot shock, and then context A with no foot shock. There was a minimum 3-hour period between exposure to context A and context B.

For the modified two-day CFD experiment, mice followed the standard CFD experiment described above for only two days, day 1 and day 2 (Extended Data Fig. 3). During these days, in context A (unsafe context), animals were placed in the conditioning chamber and allowed to freely explore for 180 seconds, after which they received a single 2 second foot shock of 1mA. Mice were taken out 15 seconds after termination of the foot shock and returned to their home cage. In context B (safe context), mice were allowed to freely explore the context for 197 seconds, the same amount of total time that they were in context A, and then returned to their home cage.

For the modified CFD experiments using stimulation in place of salient stimuli, mice placed in context A (unsafe context) were allowed to freely explore for 197 seconds, and received one of four stimulation parameters: low frequency (1-5 Hz) stimulation for 180 seconds, followed by high frequency (20-40 Hz) stimulation for 2 seconds, followed by no stimulation for 15 seconds; low frequency (1-5 Hz) stimulation for 180 seconds followed by no stimulation for 17 seconds; no stimulation for 180 seconds, followed by high frequency (20-40 Hz) stimulation for 2 seconds, followed by no stimulation for 15 seconds; no stimulation for 197 seconds. Mice were removed from the conditioning chamber after termination of the stimulation parameters and returned to their home cage. Stimulation frequencies were calibrated for each individual mouse based on fit to endogenous NE release dynamics in the CFD task during successful contextual discrimination (Fig. 1). These frequencies were tested for each mouse prior to the CFD task in a separate, neutral context.

### In Vivo Fiber Photometry

Fiber photometry recordings were made throughout the entirety of the CFD training sessions. Prior to recording, an optical fiber was attached to the implanted fiber using a ferrule sleeve (Doric, ZR_2.5). Two LEDs were used to excite GRAB_NE_ and GRAB_DA_. A 531-Hz sinusoidal LED light (Thorlabs, LED light: M470F3; LED driver: DC4104) was band-pass filtered (470 ± 20 nm, Doric, FMC4) to excite GRABNE/GRABDA and evoke NE/DA-dependent emission. A 211-Hz sinusoidal LED light (Thorlabs, LED light: M405FP1; LED driver: DC4104) was bandpass filtered (405 ± 10 nm, Doric, FMC4) to excite GRABNE/GRABDA and evoke NE/DA -independent isosbestic control emission. Prior to recording, a 300s period of GRABNE/GRABDA excitation with 405 nm and 470 nm light was used to remove the majority of baseline drift. Laser intensity for the 470 nm and 405 nm wavelength bands were measured at the tip of the optical fiber and adjusted to 50 μW before each day of recording. GRABNE/GRABDA fluorescence traveled through the same optical fiber before being bandpass filtered (525 ± 25 nm, Doric, FMC4), transduced by a femtowatt silicon photoreceiver (Newport, 2151) and recorded by a real-time processor (TDT, RZ5P). The envelopes of the 531-Hz and 211-Hz signals were extracted in real-time by the TDT program Synapse at a sampling rate of 1017.25 Hz. Optogenetic stimulation was delivered using a 625nm sinusoidal LED light (Thorlabs, LED light: M625F2; LED driver DC4104) to excite LC-DG terminals expression ChrimsonR.

### Photometry Analysis

Custom MATLAB scripts were developed for analyzing fiber photometry data in context of mouse behavior and can be accessed via GitHub (https://github.com/BruchasLab). The isosbestic 405 nm excitation control signal was subtracted from the 470 nm excitation signal to remove movement artifacts from intracellular NE-dependent GRAB_NE_/GRAB_DA_ fluorescence. Baseline drift was evident in the signal due to slow photo-bleaching artifacts, particularly during the first several minutes of each hour-long recording session. A double exponential curve was fit to the raw trace and subtracted to correct for baseline drift. After baseline correction, the photometry trace was z-scored relative to the mean and standard deviation of the test session. The post-processed fiber photometry signal was analyzed in the context of animal behavior during contextual fear discrimination task.

Quantification of NE release was obtained using the trapz function in conjunction with custom MATLAB scripts. This function utilizes numerical integration using the trapezoidal method, which approximates integration by breaking the area down into trapezoids with more easily computable areas. For an integration with N+1 evenly spaced points, the approximation is:

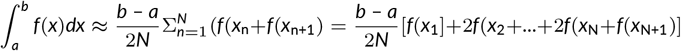

### Quantification and Statistical Analysis

All data are expressed as mean ± SEM. Behavioral data were analyzed with GraphPad Prism 10.0 (GraphPad, La Jolla, CA). Two-tailed *student’s* t-test, one-way or two-way ANOVAs were used to analyze between-subjects designs. Repeated-measures designs were analyzed using mixed-effects restricted maximum likelihood (REML) model. Tukey was used for *post-hoc* pairwise comparisons. The null hypothesis was rejected at the *p* < 0.05 level. Statistical significance was taken as ^*^*p* < 0.05, ^**^*p* < 0.01, ^***^*p* < 0.005, *n*.*s*. represents not significant. All statistical information is listed in Supplementary Table 1.

## Abbreviations

DG: dentate gyrus
LC: locus coeruleus
NE: norepinephrine.

## Materials Availability

This study did not generate new or unique reagents or other materials.

## Data Availability

Source data are provided with this paper.

## Code Availability

Custom MATLAB analysis code was created to appropriately organize, process, and combine fiber photometry data with associated behavioral data. Analysis code for photometry from Figures 1, 2, 3, and Extended Data Figures 1, 2, 3, 4, 5 will be made available online at https://www.github.com/BruchasLab.

## Acknowledgments

This work was supported by grants: National Institute of Health grants R01MH112355, F31MH130080, RF1NS118287 and P30DA048736,

National Natural Science Foundation of China grants 31925017 and 31871087, and the UW NAPE. We thank Dr. A. Suko for lab management; T. Hobbs, C. Pizzano, A. Rana, V. Lau, B. Wells for animal colony maintenance; the entire Bruchas laboratory, and multiple trainees and faculty within the NAPE Center for helpful discussions and feedback on the manuscript.

## Author Contributions

E.T.Z. performed conceptualization, methodology, investigation, formal analysis, fiber photometry analysis, writing of the original draft. G.S.S. performed behavioral experiments, immunohistochemistry. J.F. and Y.L. performed conceptualization and experiments relating to the development of the GRABNE and GRABDA sensors. M.R.B. conceptualized, acquired funding for, and supervised the project, as well as assisted in writing the manuscript.

## Lead Contact

Further information and requests for resources and reagents should be directed to and will be fulfilled by the Lead Contact, Dr. Michael Bruchas.

## SUPPLEMENTAL FIGURES

**SUPPLEMENTAL FIGURE 1.**
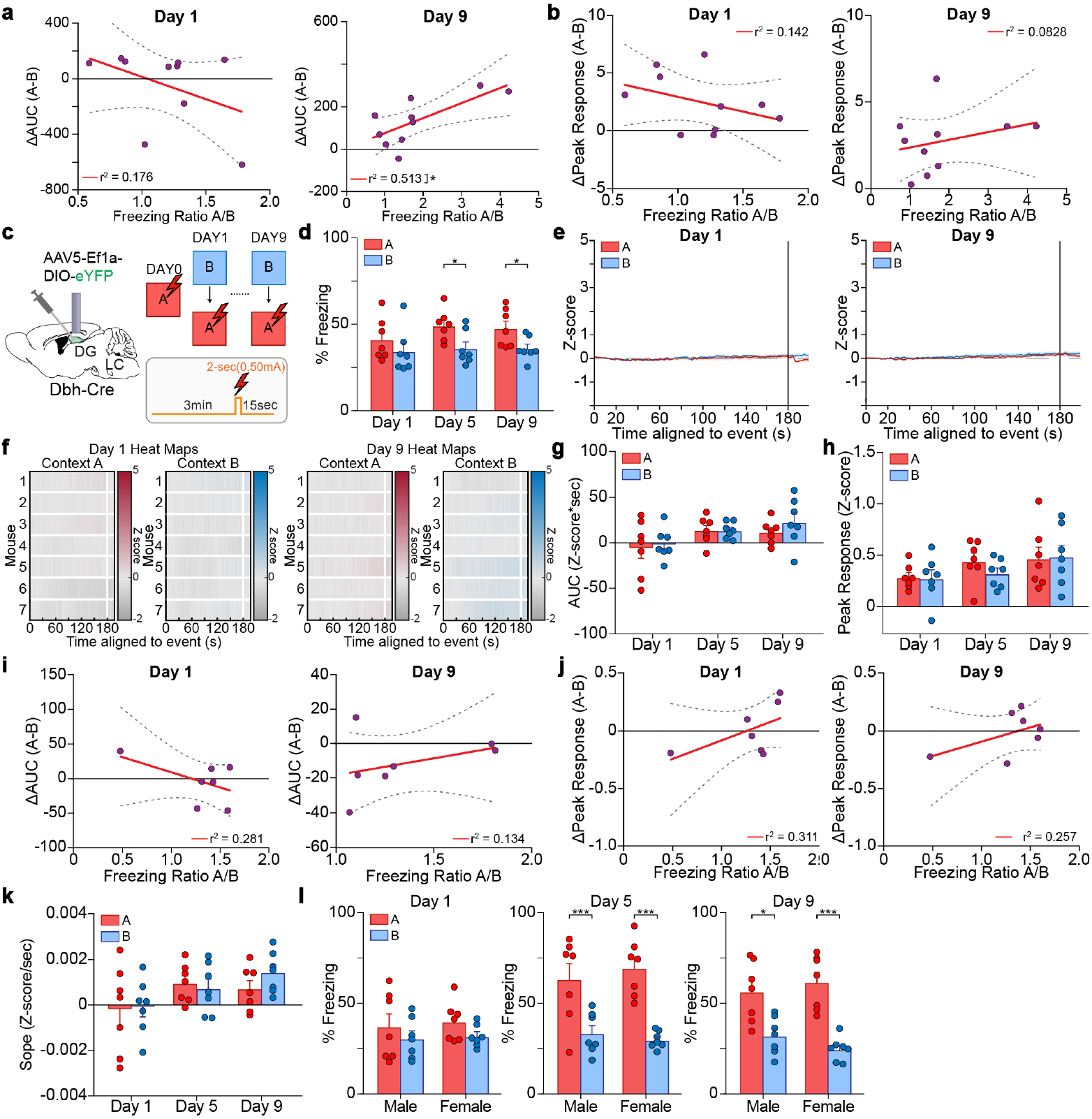
Linear elevations in NE release during contextual discrimination are detected by GRAB_NE_. Related to Fig. 1. (a) Linear regression using Pearson correlation for ratio of freezing behavior between context A and context B vs. difference in AUC between context A and context B (n = 10 mice). (b) Linear regression using Pearson correlation for ratio of freezing behavior between context A and context B vs. difference in peak response between context A and context B (n = 10 mice). (c) Schematic of experimental approach depicts infection of dentate gyrus with YFP, and optical fiber implanted above the DG (left). Schematic depicting CFD task. Animals received a shock in context A (red), but not in context B (blue) for nine days (right). (d) Mice were not able to successfully discriminate between context A (unsafe), and context B (safe) any days of training (n = 7 mice). (e) Averages traces of YFP fluorescence in context A (red) and context B (blue) for day 1 (left), and day 9 (right). (f) Individual heat maps for context A (red) and context B (blue) for days 1 (left) and 9 (right) of training. (g) Area under the curve analysis of YFP fluorescence levels in context A and context B on the first, fifth, and ninth days of training (n = 7 mice). (h) Peak response analysis of YFP fluorescence levels in context A and context B on the first, fifth, and ninth days of training (n = 7 mice). (i) Linear regression using Pearson correlation for ratio of freezing behavior between context A and context B vs. difference in AUC between context A and context B (n = 7 mice). (j) Linear regression using Pearson correlation for ratio of freezing behavior between context A and context B vs. difference in peak response between context A and context B (n = 7 mice). (k) Average slope in context A (red) and context B (blue) on the first, fifth, and ninth days of training (n = 7 mice). (L) Both male and female mice were able to successfully discriminate between context A (unsafe) and context B (safe) by the last day of training (n = 7 mice). All data are mean ± SEM. ^*^*p* < 0.05, ^***^*p* < 0.005.

**SUPPLEMENTAL FIGURE 2.**
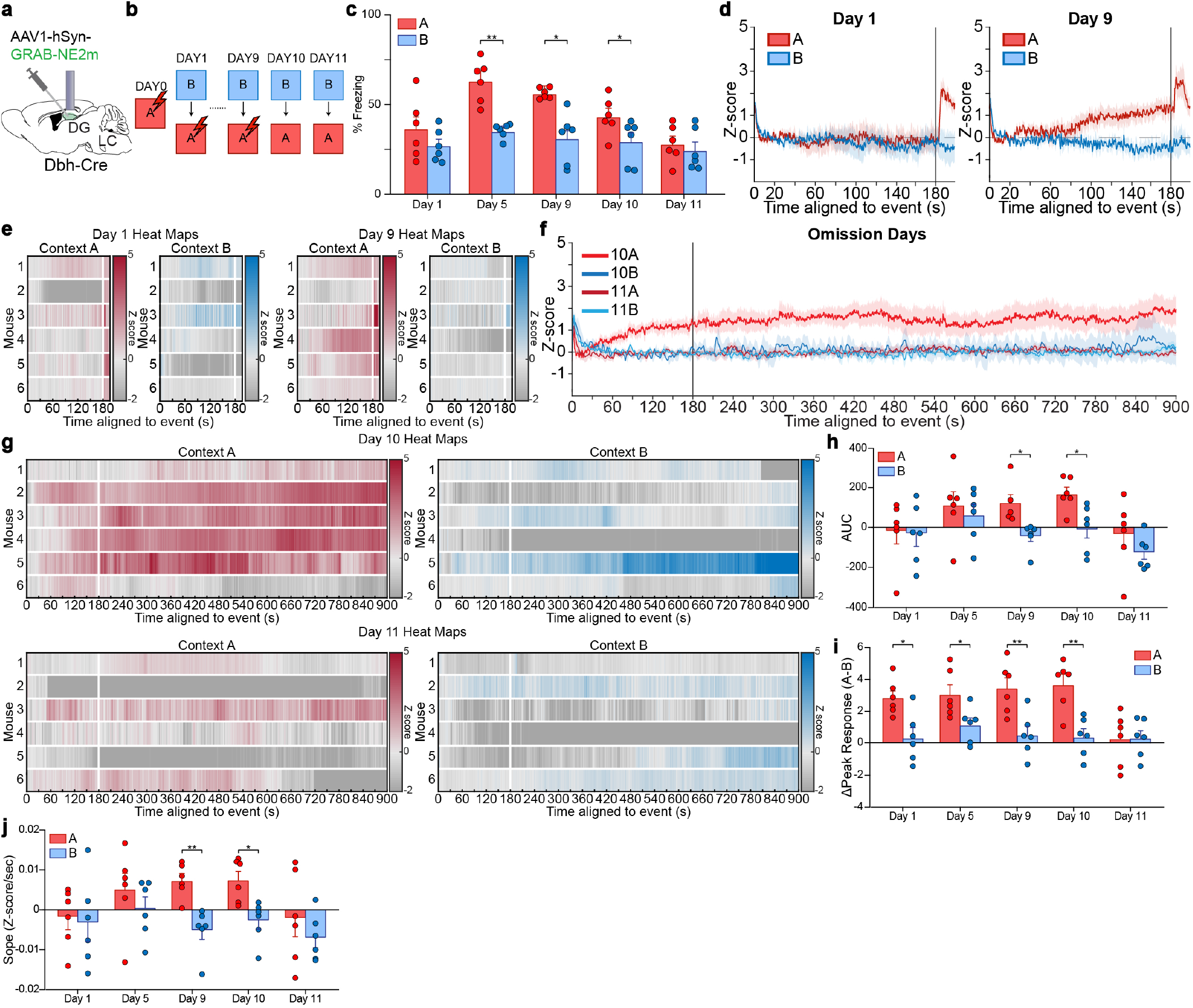
Linear elevations in NE release during contextual discrimination are extinguished following omission of salient aversive stimuli. Related to Fig. 1. (a) Schematic of experimental approach depicts infection of DG with GRAB_NE_, optical fiber implanted above the DG, and infection of LC with ChrimsonR. (b) Schematic depicting CFD task. Animals received a shock in context A (red), but not in context B (blue) for nine days. Animals then did not receive a shock in either context A (red) or context B (blue) for the following two days. (c) Mice successfully discriminated between context A (unsafe), and context B (safe) on the fifth, ninth, and tenth days of training, but not on the first and eleventh days of training (n = 6 mice). (d) Averaged traces of GRAB_NE_ fluorescence in context A (red) and context B (blue) for day 1 (left), and day 9 (right). (e) Individual heat maps for context A (red) and context B (blue) for days 1 (left) and 9 (right) of training. (f) Averaged traces of GRAB_NE_ fluorescence in context A (red) and context B (blue) for day 10 and day 11. (g) Individual heat maps for context A (red) and context B (blue) for days 10 (top) and 11 (bottom) of training. (h) Area under the curve analysis of GRAB_NE_ fluorescence levels in context A and context B on the first, fifth, ninth, tenth, and eleventh days of training (n = 6 mice). (i) Peak response analysis of GRAB_NE_ fluorescence levels in context A and context B on the first, fifth, ninth, tenth, and eleventh days of training (n = 6 mice). (j) Average slope in context A (red) and context B (blue) on the first, fifth, ninth, tenth, and eleventh days of training (n = 6 mice). All data are mean ± SEM. ^*^*p* < 0.05, ^**^*p* < 0.01.

**SUPPLEMENTAL FIGURE 3.**
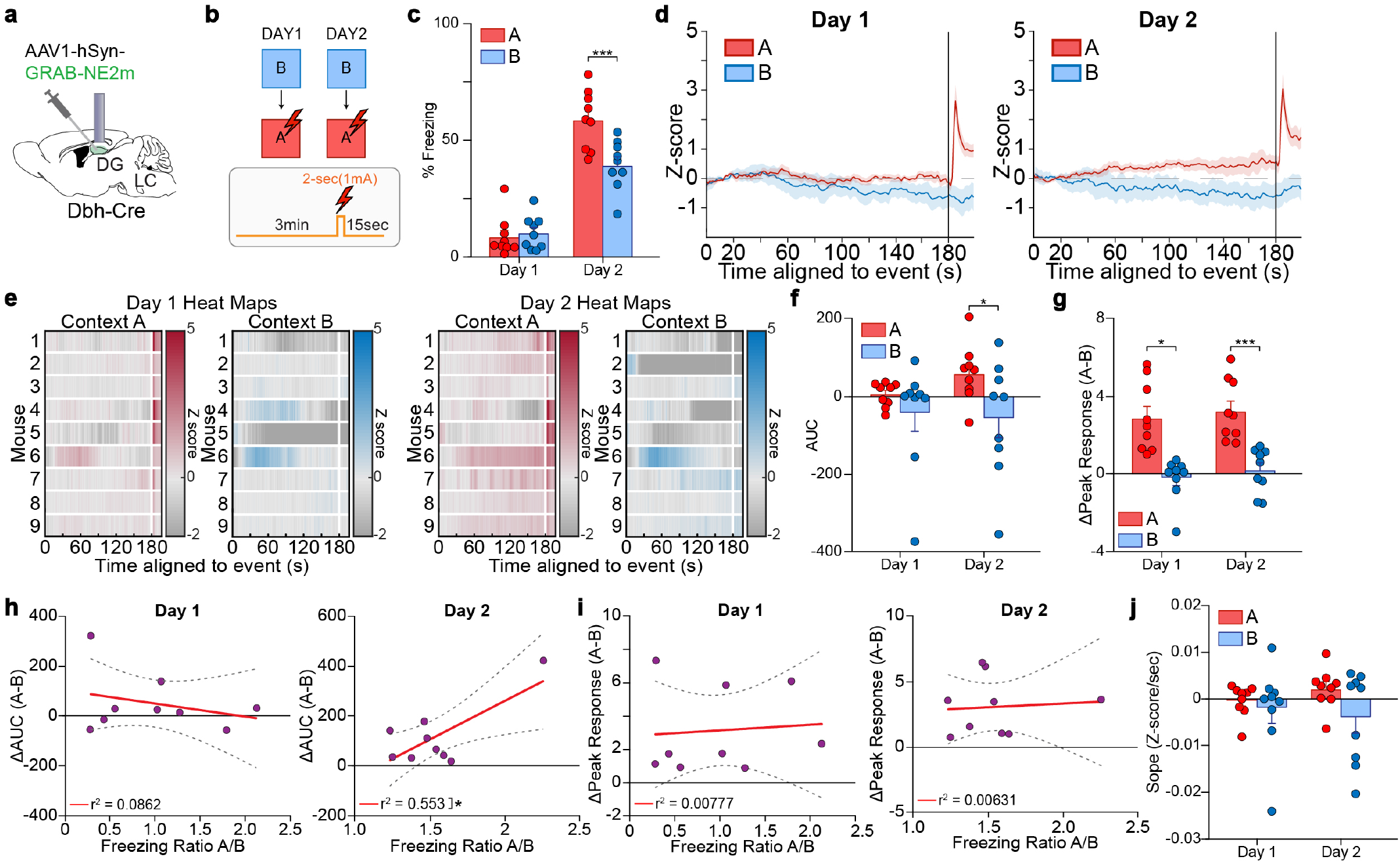
Linear elevations in NE release during contextual discrimination occur in response to a single presentation of a heightened salient aversive stimuli. Related to Fig. 1. (a) Schematic of experimental approach depicts infection of DG with GRAB_NE_, optical fiber implanted above the DG, and infection of LC with ChrimsonR. (b) Schematic depicting CFD task. Animals received a shock in context A (red), but not in context B (blue) for two days. (c) Mice successfully discriminated between context A (unsafe), and context B (safe) on the second day of training, but not on the first day of training (n = 9 mice). (d) Averaged traces of GRAB_NE_ fluorescence in context A (red) and context B (blue) for day 1 (left), and day 2 (right). (e) Individual heat maps for context A (red) and context B (blue) for days 1 (left) and 2 (right) of training. (f) Area under the curve analysis of GRAB_NE_ fluorescence levels in context A and context B on the first and second days of training (n = 9 mice). (g) Peak response analysis of GRAB_NE_ fluorescence levels in context A and context B on the first and second days of training (n = 9 mice). (h) Linear regression using Pearson correlation for ratio of freezing behavior between context A and context B vs. difference in AUC between context A and context B (n = 9 mice). (i) Linear regression using Pearson correlation for ratio of freezing behavior between context A and context B vs. difference in peak response between context A and context B (n = 9 mice). (j) Average slope in context A (red) and context B (blue) on the first and second days of training (n = 9 mice). All data are mean ± SEM. ^*^*p* < 0.05, ^***^*p* < 0.005.

**SUPPLEMENTAL FIGURE 4.**
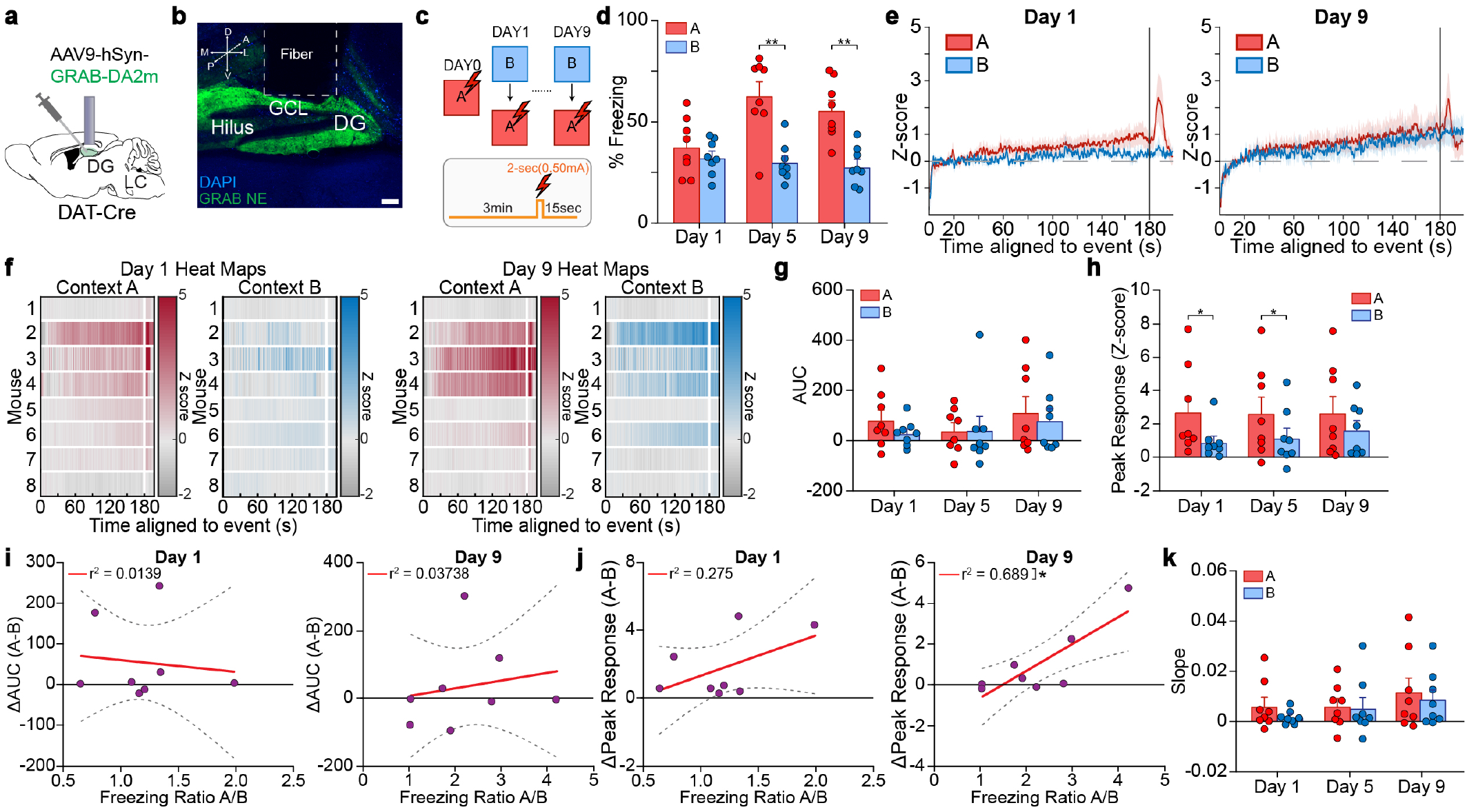
Endogenous dynamics of dopamine in the DG during contextual discrimination are detected by GRAB_DA_. Related to Fig. 1. (a) Schematic of experimental approach depicts infection of dentate gyrus with GRAB_DA_, and optical fiber implanted above the DG. (b) Representative image depicting expression of GRAB_DA_ in DG driven by hSyn promoter and location of fiber implant (n = 8 mice). Scale bar = 100 µm. (c) Schematic depicting CFD task. Animals received a shock in context A (red), but not in context B (blue) for nine days. (d) Mice successfully discriminated between context A (unsafe), and context B (safe) on the fifth and ninth days of training, but not on the first day of training (n = 8 mice). (e) Averaged traces of GRAB_DA_ fluorescence in context A (red) and context B (blue) for day 1 (left), and day 9 (right). (f) Individual heat maps for context A (red) and context B (blue) for days 1 (left) and 9 (right) of training. (g) Area under the curve analysis of GRAB_DA_ fluorescence levels in context A and context B on the first, fifth, and ninth days of training (n = 8 mice). (h) Peak response analysis of GRAB_DA_ fluorescence levels in context A and context B on the first, fifth, and ninth days of training (n = 8 mice). (i) Linear regression using Pearson correlation for ratio of freezing behavior between context A and context B vs. difference in AUC between context A and context B (n = 8 mice). (j) Linear regression using Pearson correlation for ratio of freezing behavior between context A and context B vs. difference in peak response between context A and context B (n = 8 mice). (k) Average slope in context A (red) and context B (blue) on the first, fifth, and ninth days of training (n = 8 mice). All data are mean ± SEM. ^*^*p* < 0.05, ^**^*p* < 0.01.

**SUPPLEMENTAL FIGURE 5.**
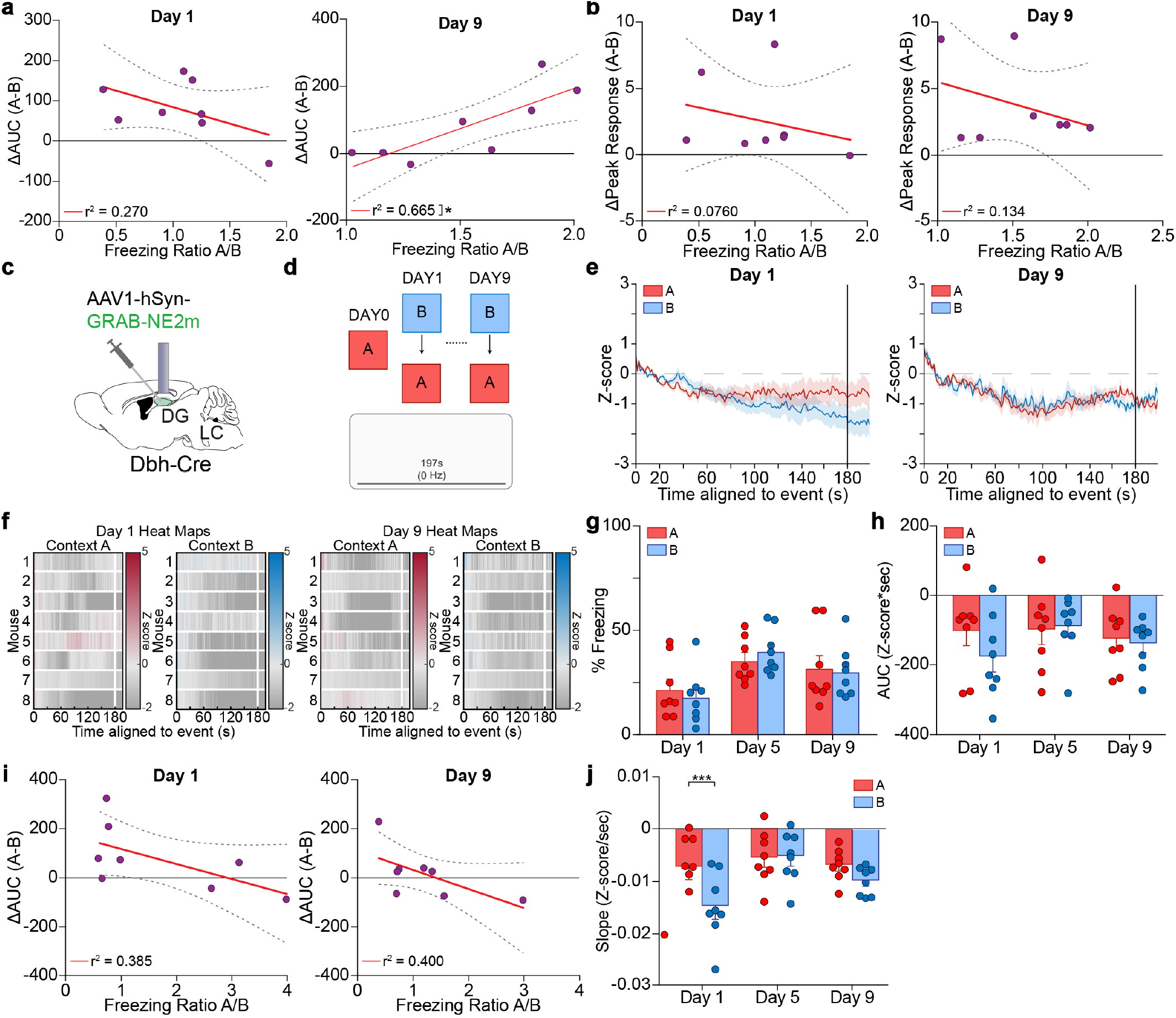
NE dynamics in the absence of a salient aversive stimuli. Related to Fig. 2. (a) Linear regression using Pearson correlation for ratio of freezing behavior between context A and context B vs. difference in AUC between context A and context B (n = 8 mice). (b) Linear regression using Pearson correlation for ratio of freezing behavior between context A and context B vs. difference in peak response between context A and context B (n = 8 mice). (c) Schematic of experimental approach depicts infection of dentate gyrus with GRAB_NE_, and optical fiber implanted above the DG. (d) Schematic depicting CFD task. Animals received no shock or stimulation in context A (red) and context B (blue) for nine days. (e) Averages traces of GRAB_NE_ fluorescence in context A (red) and context B (blue) for day 1 (left), and day 9 (right). (f) Individual heat maps for context A (red) and context B (blue) for days 1 (left) and 9 (right) of training. (g) Mice were not able to successfully discriminate between context A (unsafe), and context B (safe) any days of training (n = 8 mice). (h) Area under the curve analysis of GRAB_NE_ fluorescence levels in context A and context B on the first, fifth, and ninth days of training (n = 8 mice). (i) Linear regression using Pearson correlation for ratio of freezing behavior between context A and context B vs. difference in AUC between context A and context B (n = 8 mice). (j) Average slope in context A (red) and context B (blue) on the first, fifth, and ninth days of training (n = 8 mice). All data are mean ± SEM. ^*^*p* < 0.05, ^***^*p* < 0.005.

**SUPPLEMENTAL FIGURE 6.**
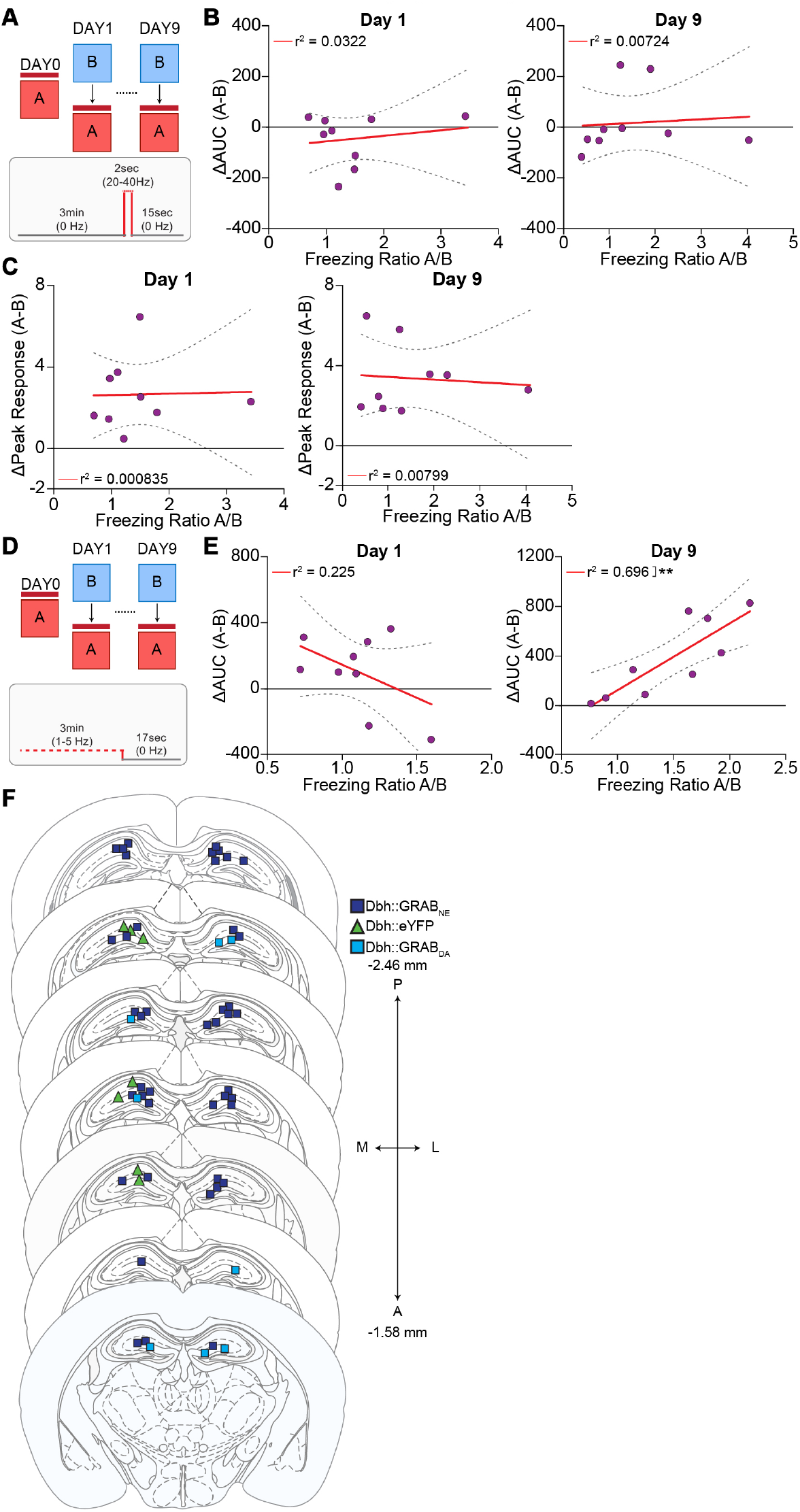
Associations between tonic and phasic stimulation in DG and contextual discrimination and optical fiber placements. Related to Fig. 1, 2, 3. (a)Schematic depicting CFD task. Animals received stimulation in context A (red), but not in context B (blue) for nine days. (b) Linear regression using Pearson correlation for ratio of freezing behavior between context A and context B vs. difference in AUC between context A and context B (n = 9 mice). (c) Linear regression using Pearson correlation for ratio of freezing behavior between context A and context B vs. difference in peak response between context A and context B. Linear regression using Pearson correlation for ratio of freezing behavior between context A and context B vs. difference in AUC between context A and context B (n = 9 mice). (d) Schematic depicting CFD task. Animals received stimulation in context A (red), but not in context B (blue) for nine days. (e) Linear regression using Pearson correlation for ratio of freezing behavior between context A and context B vs. difference in AUC between context A and context B (n = 9 mice). (f) Reconstruction of approximate locations of fiber-optic implant tips for GRAB_NE_, YFP and GRAB_DA_ mice used. All data are mean ± SEM. ^**^*p* < 0.01.

## REFERENCES

1. Liberzon I, Abelson JL. Context Processing and the Neurobiology of Post-Traumatic Stress Disorder. Neuron. 2016;92(1). Publisher: Neurondoi: 10.1016/j.neuron.2016.09.039

2. Lisman JE, Coyle JT, Green RW, et al. Circuit-based framework for understanding neurotransmitter and risk gene interactions in schizophrenia. Trends in Neurosciences. 2008;31(5):234–242. doi: 10.1016/j.tins.2008.02.005

3. Shohamy D, Mihalakos P, Chin R, Thomas B, Wagner AD, Tamminga C. Learning and generalization in schizophrenia: effects of disease and antipsychotic drug treatment. Biological Psychiatry. 2010;67(10):926–932. doi:10.1016/j.biopsych.2009.10.025

4. Kheirbek MA, Klemenhagen KC, Sahay A, Hen R. Neurogenesis and generalization: a new approach to stratify and treat anxiety disorders. Nature neuroscience. 2012;15(12). Publisher: Nat Neuroscidoi: 10.1038/nn.3262

5. Besnard A, Sahay A. Adult Hippocampal Neurogenesis, Fear Generalization, and Stress. Neuropsychopharmacology : official publication of the American College of Neuropsychopharmacology. 2016;41(1). Publisher: Neuropsychopharmacologydoi:10.1038/npp.2015.167

6. Siegmund A, Wotjak CT. Toward an Animal Model of Posttraumatic Stress Disorder. Annals of the New York Academy of Sciences. 2006;1071(1):324–334. _eprint: https://onlinelibrary.wiley.com/doi/pdf/10.1196/annals.1364.025doi:10.1196/annals.1364.025

7. Raskind MA, Peskind ER, Kanter ED, et al. Reduction of Nightmares and Other PTSD Symptoms in Combat Veterans by Prazosin: A Placebo-Controlled Study. American Journal of Psychiatry. 2003;160(2):371–373. Publisher: American Psychiatric Publishingdoi: 10.1176/appi.ajp.160.2.371

8. Taylor HR, Freeman MK, Cates ME. Prazosin for treatment of nightmares related to posttraumatic stress disorder. American Journal of Health-System Pharmacy. 2008;65(8):716–722. doi: 10.2146/ajhp070124

9. Danielson NB, Kaifosh P, Zaremba JD, et al. Distinct Contribution of Adult-Born Hippocampal Granule Cells to Context Encoding. Neuron. 2016;90(1):101. doi: 10.1016/j.neuron.2016.02.019

10. Deng W, Mayford M, Gage FH. Selection of distinct populations of dentate granule cells in response to inputs as a mechanism for pattern separation in mice. eLife. 2013;2:e00312. Publisher: eLife Sciences Publications, Ltddoi: 10.7554/eLife.00312

11. Lisman J, Buzsáki G, Eichenbaum H, Nadel L, Ranganath C, Redish AD. Viewpoints: how the hippocampus contributes to memory, navigation and cognition. Nature Neuroscience. 2017;20(11):1434–1447. Publisher: Nature Publishing Groupdoi: 10.1038/nn.4661

12. McHugh TJ, Jones MW, Quinn JJ, et al. Dentate Gyrus NMDA Receptors Mediate Rapid Pattern Separation in the Hippocampal Network. Science. 2007;317(5834):94–99. Publisher: American Association for the Advancement of Sciencedoi: 10.1126/science.1140263

13. Woods NI, Stefanini F, Apodaca-Montano DL, Tan IM, Biane JS, Kheirbek MA. The Dentate Gyrus Classifies Cortical Representations of Learned Stimuli. Neuron. 2020;107(1):173. doi: 10.1016/j.neuron.2020.04.002

14. Marr D. Simple memory: a theory for archicortex. Philosophical transactions of the Royal Society of London. Series B, Biological sciences. 1971;262(841). Publisher: Philos Trans R Soc Lond B Biol Scidoi: 10.1098/rstb.1971.0078

15. Morris RGM. Elements of a neurobiological theory of hippocampal function: the role of synaptic plasticity, synaptic tagging and schemas. European Journal of Neuroscience. 2006;23(11):2829–2846. _eprint: https://onlinelibrary.wiley.com/doi/pdf/10.1111/j.1460-9568.2006.04888.xdoi:10.1111/j.1460-9568.2006.04888.x

16. O’Reilly R, McClelland J. Hippocampal conjunctive encoding, storage, and recall: avoiding a trade-off. Hippocampus. 1994;4(6). Publisher: Hippocampusdoi: 10.1002/hipo.450040605

17. Seo Do, Zhang ET, Piantadosi SC, et al. A locus coeruleus to dentate gyrus noradrenergic circuit modulates aversive contextual processing. Neuron. 2021;109(13):2116. doi: 10.1016/j.neuron.2021.05.006

18. Britton KT, S. Segal D, Kuczenski R, Hauger R. Dissociation between in vivo hippocampal norepinephrine response and behavioral/neuroendocrine responses to noise stress in rats. Brain Research. 1992;574(1):125–130. doi: 10.1016/0006-8993(92)90808-M

19. Campeau S, Nyhuis TJ, Kryskow EM, et al. Stress rapidly increases alpha 1d adrenergic receptor mRNA in the rat dentate gyrus. Brain Research. 2010;1323:109–118. doi: 10.1016/j.brainres.2010.01.084

20. Fa M, Xia L, Anunu R, et al. Stress modulation of hippocampal activity – Spotlight on the dentate gyrus. Neurobiology of Learning and Memory. 2014;112:53–60. doi: 10.1016/j.nlm.2014.04.008

21. Blackstad T, Fuxe K, Hökfelt T. Noradrenaline nerve terminals in the hippocampal region of the rat and the guinea pig. Zeitschrift fur Zellforschung und mikroskopische Anatomie (Vienna, Austria : 1948). 1967;78(4). Publisher: Z Zellforsch Mikrosk Anatdoi: 10.1007/BF00334281

22. Ungerstedt. Stereotaxic mapping of the monoamine pathways in the rat brain. Acta physiologica Scandinavica. Supplementum. 1971;367. Publisher: Acta Physiol Scand Suppldoi: 10.1111/j.1365-201x.1971.tb10998.xbioRχiv

23. McCall JG, Al-Hasani R, Siuda ER, et al. CRH engagement of the locus coeruleus noradrenergic system mediates stress-induced anxiety. Neuron. 2015;87(3):605. doi:10.1016/j.neuron.2015.07.002

24. McCall JG, Siuda ER, Bhatti DL, et al. Locus coeruleus to basolateral amygdala noradrenergic projections promote anxiety-like behavior. eLife. 2017;6:e18247. Publisher: eLife Sciences Publications, Ltddoi: 10.7554/eLife.18247

25. Uematsu A, Tan BZ, Ycu EA, et al. Modular organization of the brainstem noradrenaline system coordinates opposing learning states. Nature Neuroscience. 2017;20(11):1602–1611. Pub-lisher: Nature Publishing Groupdoi: 10.1038/nn.4642

26. Valentino RJ, Van Bockstaele E. Convergent regulation of locus coeruleus activity as an adaptive response to stress. European Journal of Pharmacology. 2008;583(2):194–203. doi:10.1016/j.ejphar.2007.11.062

27. Harley CW. Norepinephrine and the dentate gyrus. In: Scharfman HE., ed. Progress in Brain Research,,. 163 of The Dentate Gyrus: A Comprehensive Guide to Structure, Function, and Clinical Implications. Elsevier, 2007:299–318

28. Loy R, Koziell DA, Lindsey JD, Moore RY. Noradrenergic innervation of the adult rat hippocampal formation. Journal of Comparative Neurology. 1980;189(4):699–710. _eprint: 13 42. https://onlinelibrary.wiley.com/doi/pdf/10.1002/cne.901890406doi:10.1002/cne.901890406

29. Rose GM, Pang KCH. Differential effect of norepinephrine upon granule cells and interneurons in the dentate gyrus. Brain Research. 1989;488(1):353–356. doi: 10.1016/0006-8993(89)90729-4

30. Seidenbecher T, Reymann K, Balschun D. A post-tetanic time window for the reinforcement of long-term potentiation by appetitive and aversive stimuli. Proceedings of the National Academy of Sciences. 1997;94(4):1494–1499. Publisher: Proceedings of the National Academy of Sciencesdoi: 10.1073/p-nas.94.4.1494

31. Walling SG, Harley CW. Locus Ceruleus Activation Initiates Delayed Synaptic Potentiation of Perforant Path Input to the Dentate Gyrus in Awake Rats: A Novel β-Adrenergic- and Protein Synthesis-Dependent Mammalian Plasticity Mechanism. Journal of Neuroscience. 2004;24(3):598–604. Publisher: Society for Neuroscience Section: Behavioral/Systems/Cognitivedoi:10.1523/JNEUROSCI.4426-03.2004

32. Lovett-Barron M, Kaifosh P, Kheirbek MA, et al. Dendritic Inhibition in the Hippocampus Supports Fear Learning. Science. 2014;343(6173):857–863. Publisher: American Association for the Advancement of Sciencedoi: 10.1126/science.1247485

33. Feng J, Zhang C, Lischinsky JE, et al. A genetically encoded fluorescent sensor for rapid and specific in vivo detection of norepinephrine. Neuron. 2019;102(4):745. doi:10.1016/j.neuron.2019.02.037

34. Takeuchi T, Duszkiewicz AJ, Sonneborn A, et al. Locus coeruleus and dopaminergic consolidation of everyday memory. Nature. 2016;537(7620):357–362. Publisher: Nature Publishing Groupdoi: 10.1038/nature19325

35. Kempadoo KA, Mosharov EV, Choi SJ, Sulzer D, Kandel ER. Dopamine release from the locus coeruleus to the dorsal hippocampus promotes spatial learning and memory. Proceedings of the National Academy of Sciences. 2016;113(51):14835– 14840. Publisher: Proceedings of the National Academy of Sciencesdoi: 10.1073/pnas.1616515114

36. Li L, Rana AN, Li EM, Feng J, Li Y, Bruchas MR. Activity-dependent constraints on catecholamine signaling. Cell reports. 2023;42(12):113566. doi: 10.1016/j.celrep.2023.113566

37. Howe MW, Tierney PL, Sandberg SG, Phillips PEM, Graybiel AM. Prolonged dopamine signalling in striatum signals proximity and value of distant rewards. Nature. 2013;500(7464):575–579. Publisher: Nature Publishing Groupdoi: 10.1038/nature12475

38. Lohani S, Martig AK, Underhill SM, et al. Burst activation of dopamine neurons produces prolonged post-burst availability of actively released dopamine. Neuropsychopharmacology. 2018;43(10):2083–2092. Publisher: Nature Publishing Groupdoi: 10.1038/s41386-018-0088-7

39. Wagatsuma A, Okuyama T, Sun C, Smith LM, Abe K, Tonegawa S. Locus coeruleus input to hippocampal CA3 drives singletrial learning of a novel context. Proceedings of the National Academy of Sciences. 2018;115(2):E310–E316. Publisher: Proceedings of the National Academy of Sciencesdoi: 10.1073/p-nas.1714082115

40. Brewin CR, Andrews B, Valentine JD. Meta-analysis of risk factors for posttraumatic stress disorder in trauma-exposed adults. Journal of Consulting and Clinical Psychology. 2000;68(5):748– 766. Place: US Publisher: American Psychological Associationdoi: 10.1037/0022-006X.68.5.748

41. Safari S, Ahmadi N, Mohammadkhani R, et al. Sex differences in spatial learning and memory and hippocampal long-term potentiation at perforant pathway-dentate gyrus (PP-DG) synapses in Wistar rats. Behavioral and Brain Functions. 2021;17(1):9. doi: 10.1186/s12993-021-00184-y

42. Yagi S, Lee A, Truter N, Galea LAM. Sex differences in contextual pattern separation, neurogenesis, and functional connectivity within the limbic system. Biology of Sex Differences. 2022;13(1):42. doi: 10.1186/s13293-022-00450-2

43. Bonanno GA, Romero SA, Klein SI. The temporal elements of psychological resilience: An integrative framework for the study of individuals, families, and communities. Psychological Inquiry. 2015;26(2):139–169. Place: United Kingdom Publisher: Taylor & Francisdoi: 10.1080/1047840X.2015.992677

44. Kessler RC, Sonnega A, Bromet E, Hughes M, Nelson CB. Posttraumatic Stress Disorder in the National Comorbidity Survey. Archives of General Psychiatry. 1995;52(12):1048–1060. doi: 10.1001/archpsyc.1995.03950240066012

45. Hunsaker MR, Rosenberg JS, Kesner RP. The role of the dentate gyrus, CA3a,b, and CA3c for detecting spatial and environmental novelty. Hippocampus. 2008;18(10):1064–1073. _eprint: https://onlinelibrary.wiley.com/doi/pdf/10.1002/hipo.20464doi:10.1002/hipo.20464

46. Prince LY, Bacon TJ, Tigaret CM, Mellor JR. Neuromodulation of the Feedforward Dentate Gyrus-CA3 Microcircuit. Frontiers in Synaptic Neuroscience. 2016;8. Publisher: Frontiersdoi: 10.3389/fnsyn.2016.00032

