## Supplementary Table 1 for "Dentate Gyrus Norepinephrine Ramping Facilitates Aversive Contextual Processing"

| Figure | | Experimental Variables | Statistical Test | Results |
|---|---|---|---|---|
| 1 | e | Behavior (Tail lift) | Unpaired t-test | GRAB-NE vs YFP: $t_{(13)} = 4.35$; p = 0.0008 |
| | g | Behavior (Foot shock) | Unpaired t-test | GRAB-NE vs YFP: $t_{(12)} = 3.502$; p = 0.0044 |
| | k | Behavior (CFD) | Paired t-test | Day 1: $t_{(9)} = 2.048$, p = 0.0709 |
| | | | | Day 5: $t_{(9)} = 4.137$, p = 0.0025 |
| | | | | Day 9: $t_{(9)} = 2.660$, p = 0.0261 |
| | l | Area Under Curve | Paired t-test | Day 1: $t_{(9)} = 0.519$, p = 0.6163 |
| | | | | Day 5: $t_{(9)} = 3.355$, p = 0.0085 |
| | | | | Day 9: $t_{(9)} = 3.774$, p = 0.0044 |
| | m | Peak Response | Paired t-test | Day 1: $t_{(9)} = 3.111$, p = 0.0125 |
| | | | | Day 5: $t_{(9)} = 5.316$, p = 0.0005 |
| | | | | Day 9: $t_{(9)} = 4.931$, p = 0.0008 |
| | n | Slope | Paired t-test | Day 1: $t_{(9)} = 1.160$, p = 0.2758 |
| | | | | Day 5: $t_{(9)} = 2.586$, p = 0.0294 |
| | | | | Day 9: $t_{(9)} = 3.870$, p = 0.0038 |
| 2 | e | Optogenetic stimulation (ChrimsonR) | Unpaired t-test | GRAB-NE vs YFP: $t_{(13)} = 3.859$; p = 0.002 |
| | h | Optogenetic stimulation (ChrimsonR) | Tukey's multiple comparisons | 1 Hz vs. 3 Hz: p = 0.7718 |
| | | | | 1 Hz vs. 5 Hz: p = 0.0007 |
| | | | | 1 Hz vs. 10 Hz: p < 0.0001 |
| | | | | 1 Hz vs. 20 Hz: p < 0.0001 |
| | | | | 3 Hz vs. 5 Hz: p = 0.0177 |
| | | | | 3 Hz vs. 10 Hz: p < 0.0001 |
| | | | | 3 Hz vs. 20 Hz: p < 0.0001 |
| | | | | 5 Hz vs. 10 Hz: p = 0.0466 |
| | | | | 5 Hz vs. 20 Hz: p < 0.0001 |
| | | | | 10 Hz vs. 20 Hz: p = 0.0005 |
| | l | Behavior (CFD) | Paired t-test | Day 1: $t_{(7)} = 0.4134$, p = 0.6917 |
| | | | | Day 5: $t_{(7)} = 1.998$, p = 0.0859 |
| | | | | Day 9: $t_{(7)} = 4.184$, p = 0.0041 |
| | m | Area Under Curve | Paired t-test | Day 1: $t_{(7)} = 3.111$, p = 0.0170 |
| | | | | Day 5: $t_{(7)} = 3.033$, p = 0.0190 |
| | | | | Day 9: $t_{(7)} = 4.025$, p = 0.005 |
| | n | Peak Response | Paired t-test | Day 1: $t_{(7)} = 2.390$, p = 0.0482 |
| | | | | Day 5: $t_{(7)} = 2.446$, p = 0.0443 |
| | | | | Day 9: $t_{(7)} = 3.260$, p = 0.0139 |
| | o | Slope | Paired t-test | Day 1: $t_{(7)} = 1.538$, p = 0.168 |
| | | | | Day 5: $t_{(7)} = 3.661$, p = 0.0081 |
| | | | | Day 9: $t_{(7)} = 4.039$, p = 0.0049 |

| | | | | |
|---|---|---|---|---|
| 3 | f | Behavior (CFD) | Paired t-test | Day 1: $t_{(8)}$ = 1.480, p = 0.1773 |
| | | | | Day 5: $t_{(8)}$ = 1.114, p = 0.2975 |
| | | | | Day 9: $t_{(8)}$ = 0.9306, p = 0.3793 |
| | g | Area Under Curve | Paired t-test | Day 1: $t_{(8)}$ = 1.344, p = 0.2158 |
| | | | | Day 5: $t_{(8)}$ = 1.090, p = 0,3074 |
| | | | | Day 9: $t_{(8)}$ = 0.4070, p = 0.6947 |
| | h | Peak Response | Paired t-test | Day 1: $t_{(7)}$ = 4.519, p = 0.0020 |
| | | | | Day 5: $t_{(7)}$ = 3.535, p = 0.0077 |
| | | | | Day 9: $t_{(7)}$ = 5.824, p = 0.0004 |
| | h | Slope | Paired t-test | Day 1: $t_{(7)}$ = 1.380, p = 0.2051 |
| | | | | Day 5: $t_{(7)}$ = 1.085, p = 0.3097 |
| | | | | Day 9: $t_{(7)}$ = 0.8658, p = 0.4118 |
| | o | Behavior (CFD) | Paired t-test | Day 1: $t_{(8)}$ = 0.9955, p = 0.3486 |
| | | | | Day 5: $t_{(8)}$ = 2.072, p = 0.0720 |
| | | | | Day 9: $t_{(8)}$ = 2.735, p = 0.0257 |
| | p | Area Under Curve | Paired t-test | Day 1: $t_{(8)}$ = 1.335, p = 0.2187 |
| | | | | Day 5: $t_{(8)}$ = 5.359, p = 0.0007 |
| | | | | Day 9: $t_{(8)}$ = 3.611, p = 0.0069 |
| | q | Slope | Paired t-test | Day 1: $t_{(8)}$ = 3.197, p = 0.0127 |
| | | | | Day 5: $t_{(8)}$ = 4.235, p = 0.0029 |
| | | | | Day 9: $t_{(8)}$ = 3.053, p = 0.0157 |
| S1 | a | Freezing Response-ΔAUC Correlation | Pearson correlation | Day 1: r = -0.4196; $r^2$ = 0.176; p = 0.2274 |
| | | | | Day 9: r = 0.7164; $r^2$ = 0.5133; p = 0.0198 |
| | b | Freezing Response-ΔPeak Response Correlation | Pearson correlation | Day 1: r = -0.3773; $r^2$ = 0.1424; p = 0.2824 |
| | | | | Day 9: r = 0.2878; $r^2$ = 0.08281; p = 0.4201 |
| | d | Behavior (YFP) | Paired t-test | Day 1: $t_{(6)}$ = 1.034, p = 0.3411 |
| | | | | Day 5: $t_{(6)}$ = 2.610, p = 0.0401 |
| | | | | Day 9: $t_{(6)}$ = 3.197, p = 0.0187 |
| | g | Area Under Curve | Paired t-test | Day 1: $t_{(6)}$ = 0.3533, p = 0.7359 |
| | | | | Day 5: $t_{(6)}$ = 0.05449, p = 0.9583 |
| | | | | Day 9: $t_{(6)}$ = 1.722, p = 0.1358 |
| | h | Peak Response | Paired t-test | Day 1: $t_{(6)}$ = 0.1490, p = 0.8865 |
| | | | | Day 5: $t_{(6)}$ = 1.248, p = 0.2586 |
| | | | | Day 9: $t_{(6)}$ = 0.2807, p = 0.7884 |
| | i | Freezing Response-ΔAUC Correlation | Pearson correlation | Day 1: r = -0.5304; $r^2$ = 0.2813; p = 0.2207 |
| | | | | Day 9: r = 0.3660; $r^2$ = 0.1339; p = 0.4195 |
| | j | Freezing Response-ΔPeak Response Correlation | Pearson correlation | Day 1: r = 0.5576; $r^2$ = 0.3109; p = 0.1934 |
| | | | | Day 9: r = 0.5066; $r^2$ = 0.2566; p = 0.2460 |

| | | | | |
|---|---|---|---|---|
| | k | Slope | Paired t-test | Day 1: $t_{(6)}$ = 0.1646, p = 0.8747 |
| | | | | Day 5: $t_{(6)}$ = 0.4062, p = 0.6987 |
| | | | | Day 9: $t_{(6)}$ = 2.178, p = 0.0722 |
| | l | Behavior (Male vs. Female) | Tukey's multiple comparisons | Male 1A vs. 1B: p = 0.9334 |
| | | | | Male 5A vs. 5B p = 0.0007 |
| | | | | Male 9A vs. 9B = 0.0110 |
| | | | | Female 1A vs. 1B = 0.8501 |
| | | | | Female 5A vs. 5B < 0.0001 |
| | | | | Female 9A vs. 9B < 0.0001 |
| | | | | Male 1A vs. Female 1A: p = 0.9986 |
| | | | | Male 1B vs. Female 1B: p > 0.9999 |
| | | | | Male 5A vs. Female 5A: p = 0.9469 |
| | | | | Male 5B vs. Female 5B: p = 0.9961 |
| | | | | Male 9A vs. Female 9A: p = 0.9753 |
| | | | | Male 9B vs. Female 9B: p = 0.9107 |
| S2 | c | Behavior (CFD) | Paired t-test | Day 1: $t_{(5)}$ = 2.563, p = 0.0505 |
| | | | | Day 5: $t_{(5)}$ = 4.938, p = 0.0043 |
| | | | | Day 9: $t_{(5)}$ = 3.926, p = 0.0111 |
| | | | | Day 10: $t_{(5)}$ = 4.229, p = 0.0083 |
| | | | | Day 11: $t_{(5)}$ = 0.06208, p = 0.9529 |
| | h | Area Under Curve | Paired t-test | Day 1: $t_{(5)}$ = 0.2734, p = 0.7955 |
| | | | | Day 5: $t_{(5)}$ = 1.192, p = 0.2866 |
| | | | | Day 9: $t_{(5)}$ = 2.600, p = 0.0482 |
| | | | | Day 10: $t_{(5)}$ = 2.598, p = 0.0484 |
| | | | | Day 11: $t_{(5)}$ = 1.060, p = 0.3377 |
| | i | Peak Response | Paired t-test | Day 1: $t_{(5)}$ = 3.950, p = 0.0108 |
| | | | | Day 5: $t_{(5)}$ = 3.858, p = 0.0119 |
| | | | | Day 9: $t_{(5)}$ = 4.431, p = 0.0068 |
| | | | | Day 10: $t_{(5)}$ = 4.229, p = 0.0083 |
| | | | | Day 11: $t_{(5)}$ = 0.06208, p = 0.9529 |
| | j | Slope | Paired t-test | Day 1: $t_{(5)}$ = 0.4971, p = 0.6402 |
| | | | | Day 5: $t_{(5)}$ = 1.652, p = 0.1595 |
| | | | | Day 9: $t_{(5)}$ = 4.660, p = 0.0055 |
| | | | | Day 10: $t_{(5)}$ = 2.738, p = 0.0409 |
| | | | | Day 11: $t_{(5)}$ = 0.8446, p = 0.4369 |
| S3 | c | Behavior (CFD) | Paired t-test | Day 1: $t_{(8)}$ = 0.6063, p = 0.5611 |
| | | | | Day 2: $t_{(8)}$ = 9.136, p < 0.0001 |
| | f | Area Under Curve | Paired t-test | Day 1: $t_{(8)}$ = 1.254, p = 0.2453 |
| | | | | Day 2: $t_{(8)}$ = 2.744, p = 0.0253 |
| | g | Peak Response | Paired t-test | Day 1: $t_{(8)}$ = 3.649, p = 0.0065 |
| | | | | Day 2: $t_{(8)}$ = 4.201, p = 0.0030 |
| | h | Freezing Response-ΔAUC Correlation | Pearson correlation | Day 1: r = -0.2936; $r^2$ = 0.08622; p = 0.4432 |
| | | | | Day 9: r = 0.7437; $r^2$ = 0.5530; p = 0.0216 |
| | i | Freezing Response-ΔPeak Response Correlation | Pearson correlation | Day 1: r = 0.08815; $r^2$ = 0.00777; p = 0.8216 |

| | | | | |
|---|---|---|---|---|
| | | | | Day 9: $r = 0.07946$; $r^2 = 0.006314$; $p = 0.8390$ |
| | j | Slope | Paired t-test | Day 1: $t_{(8)} = 0.6425$, $p = 0.5385$ |
| | | | | Day 2: $t_{(8)} = 2.081$, $p = 0.0710$ |
| S4 | d | Behavior (CFD) | Paired t-test | Day 1: $t_{(7)} = 1.174$, $p = 0.2788$ |
| | | | | Day 5: $t_{(7)} = 5.208$, $p = 0.0012$ |
| | | | | Day 9: $t_{(7)} = 3.962$, $p = 0.0055$ |
| | g | Area Under Curve | Paired t-test | Day 1: $t_{(7)} = 1.553$, $p = 0.1643$ |
| | | | | Day 5: $t_{(7)} = 0.0007542$, $p = 0.9994$ |
| | | | | Day 9: $t_{(7)} = 0.7761$, $p = 0.4631$ |
| | h | Peak Response | Paired t-test | Day 1: $t_{(7)} = 2.694$, $p = 0.0309$ |
| | | | | Day 5: $t_{(7)} = 2.482$, $p = 0.0421$ |
| | | | | Day 9: $t_{(7)} = 1.675$, $p = 0.1379$ |
| | i | Freezing Response-ΔAUC Correlation | Pearson correlation | Day 1: $r = -0.1179$; $r^2 = 0.0139$; $p = 0.7810$ |
| | | | | Day 9: $r = 0.1933$; $r^2 = 0.03738$; $p = 0.6464$ |
| | j | Freezing Response-ΔPeak Response Correlation | Pearson correlation | Day 1: $r = 0.5241$; $r^2 = 0.2746$; $p = 0.1825$ |
| | | | | Day 9: $r = 0.8301$; $r^2 = 0.6891$; $p = 0.0108$ |
| | k | Slope | Paired t-test | Day 1: $t_{(7)} = 1.285$, $p = 0.2396$ |
| | | | | Day 5: $t_{(7)} = 0.1914$, $p = 0.8537$ |
| | | | | Day 9: $t_{(7)} = 0.6831$, $p = 0.5165$ |
| S5 | a | Freezing Response-ΔAUC Correlation | Pearson correlation | Day 1: $r = -0.5197$; $r^2 = 0.2701$; $p = 0.1868$ |
| | | | | Day 9: $r = 0.8152$; $r^2 = 0.6645$; $p = 0.0137$ |
| | b | Freezing Response-ΔPeak Response Correlation | Pearson correlation | Day 1: $r = -0.2756$; $r^2 = 0.07596$; $p = 0.5088$ |
| | | | | Day 9: $r = -0.3664$; $r^2 = 0.1343$; $p = 0.3720$ |
| | g | Behavior (Control) | Paired t-test | Day 1: $t_{(7)} = 0.7717$, $p = 0.4655$ |
| | | | | Day 5: $t_{(7)} = 2.130$, $p = 0.0707$ |
| | | | | Day 9: $t_{(7)} = 0.2125$, $p = 0.8378$ |
| | h | Area Under Curve | Paired t-test | Day 1: $t_{(7)} = 1.573$, $p = 0.1598$ |
| | | | | Day 5: $t_{(7)} = 0.2512$, $p = 0.8089$ |
| | | | | Day 9: $t_{(7)} = 0.4359$, $p = 0.6760$ |
| | i | Freezing Response-ΔAUC Correlation | Pearson correlation | Day 1: $r = -0.6201$; $r^2 = 0.3846$; $p = 0.1010$ |
| | | | | Day 9: $r = -0.6320$; $r^2 = 0.3995$; $p = 0.0927$ |
| | j | Slope | Paired t-test | Day 1: $t_{(7)} = 6.560$, $p = 0.0003$ |
| | | | | Day 5: $t_{(7)} = 0.05096$, $p = 0.9608$ |
| | | | | Day 9: $t_{(7)} = 1.830$, $p = 0.1099$ |
| S6 | b | Freezing Response-ΔAUC Correlation | Pearson correlation | Day 1: $r = 0.1793$; $r^2 = 0.03215$; $p = 0.6444$ |
| | | | | Day 9: $r = 0.08510$; $r^2 = 0.007241$; $p = 0.8277$ |
| | c | Freezing Response-ΔPeak Response Correlation | Pearson correlation | Day 1: $r = 0.02890$; $r^2 = 0.0008350$; $p = 0.9412$ |
| | | | | Day 9: $r = -0.08937$; $r^2 = 0.007987$; $p = 0.8191$ |

| | | | | |
|---|---|---|---|---|
| | d | Freezing Response-ΔAUC Correlation | Pearson correlation | Day 1: r = -0.4744; $r^2$ = 0.2250; p = 0.1970 |
| | | | | Day 9: r = 0.8344; $r^2$ = 0.6963; p = 0.0052 |
